# Photosynthesis regulation impacts carbon and nitrogen assimilation in the diazotrophic cyanobacterium *Anabaena* sp. PCC 7120

**DOI:** 10.1101/2025.04.16.649060

**Authors:** Anna Santin, Mattia Storti, Massimo Mezzavilla, Anna Fortunato, Francesca Arcudi, Enrique Flores, Tomas Morosinotto, Giorgio Perin

**Affiliations:** Department of Biology, University of Padova, 35131, Padova, Italy; Department of Chemistry, University of Padova, 35131, Padova, Italy; Instituto de Bioquímica Vegetal y Fotosíntesis, CSIC and University of Seville, Seville, Spain

**Keywords:** *Anabaena* sp. PCC 7120, Cyanobacteria, Electron transport, Metabolism, Nitrogen fixation, Photosynthesis

## Abstract

- Diazotrophic cyanobacteria fix both atmospheric carbon (C) and nitrogen (N) into biomass, but the two assimilation pathways are not compatible. Species like *Anabaena* sp. PCC 7120 physically separates C and N assimilation in different cell types. Even if separated, they are strongly intertwined, as N assimilation relies on the C skeletons and reducing power from photosynthesis, that in turn depends on N rich molecules as pigments and proteins.
- Whereas the two pathways have been extensively studied individually, here we investigate their interaction by analysing photosynthetic properties upon exposure to changes in light, CO_2_ and N availability, including the contribution of photosynthetic electron fluxes.
- Growth depended on the availability of both light and CO_2_, while the N_2_ fixation activity mainly on the C supply. Upon diazotrophic conditions, the total photosynthetic electron transport activity increased, with a modified contribution of different electron pathways. A mutant strain affected in the vehiculation of fixed N between cell types showed that the modulation of photosynthesis depended on the metabolic connection between assimilation pathways.
- Overall, data showed that the regulation of photosynthetic electron fluxes is a major component of the synergic metabolic relationship between C and N assimilation pathways upon dynamic environmental conditions.

## Introduction

Cyanobacteria are a group of diverse prokaryotic organisms able to perform oxygenic photosynthesis (Singh & Montgomery, 2011), thus the conversion of carbon dioxide (CO_2_) into biomass through the exploitation of sunlight energy (Kolber *et al*., 2001; Fowler *et al*., 2013). They are widespread in many different habitats where they are major contributors to net primary productivity (Mangan *et al*., 2016; Lee *et al*., 2021).

Some cyanobacteria, called diazotrophs, are also capable of fixing atmospheric nitrogen (N_2_) into biologically assimilable chemical species, such as ammonium (NH_4_^+^), in a process called Biological N Fixation (BNF). BNF is performed only by some prokaryotes (i.e. archaea and bacteria including cyanobacteria) and is estimated to enable globally the fixation of approximately 240 Tg N per year with an essential contribution to the N cycle (Issa *et al*., 2014). Diazotrophic cyanobacteria thus fix both atmospheric carbon (C) and N into biomass, placing them at a strategic intersection between the biogeochemical cycles of these two elements in nature, where they contribute to the bioavailability of essential chemical species to sustain the whole biosphere.

Nitrogenases, the enzymes responsible for BNF, are oxygen (O_2_) sensitive (Fay & Cox, 1967; Wong & Burris, 1972) and N_2_ fixation is therefore not biologically compatible with the photosynthetic generation of O_2_. Diazotrophic cyanobacteria bypassed this limitation by evolving strategies for either a temporal or spatial segregation between the two processes (Berman-Frank *et al*., 2001). As example of the latter, in the absence of combined N sources (e.g. NO_3_^-^ or NH_4_^+^) in the medium, filamentous cyanobacteria, such as the model *Anabaena* sp. PCC 7120, differentiate 5-10% of their photosynthetically active vegetative cells (VCs) into heterocysts (HCs) (Kumar *et al*., 2010), where N_2_ fixation occurs (Elhai & Wolk, 1990; Berman-Frank *et al*., 2001; Herrero *et al*., 2016).

To maintain a microoxic environment in HCs, photosystem II (PSII) is dismantled to prevent O_2_ evolution and an extensive metabolic remodelling takes place (Golden & Yoon, 1998), including Rubisco degradation and increased O_2_ scavenging activity, catalysed by both flavodiiron proteins (Ermakova *et al*., 2014) and respiration (Valladares *et al*., 2007). Moreover, HCs undergo a structural remodelling, through the deposition of two additional layers to their cell wall, one of which is a glycolipid layer that represents a hydrophobic barrier to O_2_, thus preventing its entrance by diffusion (Nicolaisen *et al*., 2009; Shvarev *et al*., 2019).

Under these conditions, cyanobacterial filaments thus include two different cell types, HCs and VCs, with specialised metabolisms, respectively dedicated to sustaining N and C assimilation pathways. However, N assimilation depends on the availability of C skeletons generated using energy and reducing power provided by photosynthesis, whilst photosynthetic functionality depends on the availability of pigments and proteins synthesized thanks to N assimilation, making the two pathways strongly intertwined. The metabolic complementarity of HCs and VCs (Nieves-Morión *et al*., 2021) calls for a tight coordination to ensure whole-system homeostasis (Zhang *et al*., 2018). In fact, such robust connection between the two assimilation pathways is needed also to maintain a strict C/N ratio, which upon balanced growth conditions stands at approximately 5:1 (Wolk, 1973; Forchhammer & Selim, 2019). This is likely to depend on a complex network of sensors, metabolic effectors and protein targets (Herrero & Flores, 2019; Forchhammer & Selim, 2019) whose impact on the overall process performances is still poorly known. Moreover, in nature, this coordination needs also great versatility to properly tune the allocation of resources between metabolic pathways and regulate growth as a function of dynamic environmental inputs (e.g. light, nutrients and CO_2_) (Perin *et al*., 2021).

The objective of this work was to investigate the impact of photosynthesis regulation on the cooperation between C and N assimilation pathways in *Anabaena* sp. PCC 7120 in response to variable metabolic inputs, resembling the natural dynamics. Systematic variations of irradiance, CO_2_ and N availability were exploited to alter both the C and N assimilation pathways and revealed that photosynthetic activity responds to the condition of N_2_ fixation. Data from the wild-type strain were compared to a mutant affected in the vehiculation of fixed N from HCs to VCs (Burnat *et al*., 2014), revealing both that the functional metabolic dependence between cell types is reciprocal and that photosynthesis regulation is a relevant component of the metabolic coordination between C and N assimilation pathways, likely to optimize the allocation of resources between cells types under dynamic environmental conditions.

## Material and Methods

### Growth conditions

Axenic *Anabaena* sp. (also known as *Nostoc* sp.) PCC 7120 (hereafter *Anabaena*) cultures were maintained in BG11 media (Rippka *et al*., 1979) with continuous shaking at 90 rpm, 30 µmol photons m^-2^ s^-1^ (US-SQS/L, Walz, Germany) illumination by white 3000 K LED source (C-LED, Italy) and 30°C.

For experiments, 20 mL of *Anabaena* cultures were washed twice in the final cultivation medium and grown in 100 mL Erlenmeyer flasks for 4 days starting from an initial optical density at 750 nm (OD_750_) of 0.05 (Spark, Tecan, Switzerland). BG11 and BG11_0_ (BG11 depleted in NaNO_3_) were chosen as different N conditions; 30 and 300 µmol photons m^-2^ s^-1^ were set as different light intensities using a LI-250A luminometer (Heinz-Waltz, Effeltrich, Germany), provided by 3000 K LED source (C-LED, Italy); air (350-430 ppm CO_2_) or enriched CO_2_ (10000 ppm CO_2_), maintained through a control system (EcoTechnics Evolution, Servovendi, Italy), were used as different C supply. The growth rate was calculated from daily OD_750_ measurements:

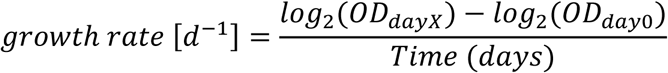

where X is the last day of the exponential growth phase. For dry weight determinations, 10 mL of cultures were filtered on 0.45 µm Cellulose Nitrate Filter (Sartorius, Germany), according to Perin et al. (2015).

### Pigments determination

*Anabaena* cultures were pelleted at 18000 g for 15 minutes and resuspended in an iso-volume (1 mL) of 100% methanol, then incubated in the dark at 4°C for 2 hours. Insoluble debris was pelleted at 18000 g for 15 minutes, and absorption of 200 µL of supernatant was measured in technical duplicate with a multi-well plates spectrophotometer (Spark, Tecan, Switzerland). Carotenoids and chlorophyll a (Chl) content was quantified from the absorption values at 470, 665 and 730 nm, according to Ritchie (2006):

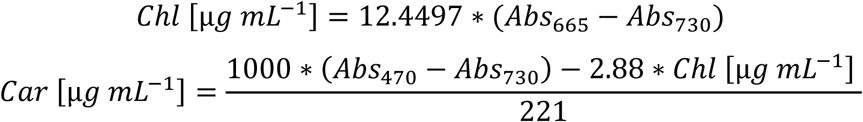

### Photosynthetic activity

Spectroscopic analysis of P700 in Photosystem I (PSI) was performed with a JTS-10 spectrophotometer (Biologic, France) similarly to Simionato et al. (2013) (Supplementary Fig. S1a for a detailed description of the method). Briefly, 4 days-old cultures of *Anabaena* were resuspended in either fresh BG11 or BG11_0_ medium, depending on the experimental conditions, at a concentration of 5 µg mL^-1^ Chl. Upon dark exposure, PSI is fully reduced (Supplementary Fig. S1a). Cyanobacteria were illuminated for 15 seconds by an orange LED ring (630 nm) to drive PSI oxidation (P700^+^) and the relaxation of the signal was followed for 10 seconds in the dark. The experiment was reiterated in presence of either 20 µM DCMU, to block the PSII-dependent electron flow, or 20 µM DCMU + 40 µM DBIMB to block the electron flow through cytochrome *b_6_f* (Cyt *b_6_f*). These concentrations are the result of preliminary investigations to identify the minimum value enabling maximal PSI oxidation upon illumination whilst avoiding side effects like photoinhibition (Supplementary Figure S2). The signal was measured with a detection light (700 nm, FWHM 25 nm) passing through a 705 nm (FWHM 6 nm) interference filter and a not-specific wavelength depleted by a P700-705 cut-off filter (705 nm, FWHM 6 nm, Biologic, France).

Maximum P700 oxidation (P700_max_) and relative abundance was measured by the 705 nm maximum absorption light-dark difference. The Total Electron Flow (TEF) was calculated from the Oxidation rate (P700^+^/P700) and the time needed for P700 reduction at light offset (t_2/3_, Reduction Rate), according to the following equation:

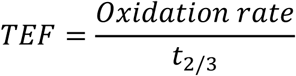

Without inhibitors, linear (LEF), cyclic (CEF) and alternative (AEF) electron flows are active. In this condition, they altogether contribute to the total electron flow (TEF), with TEF = LEF + CEF + AEF. In presence of DCMU, LEF is instead inactivated, with TEF_DCMU_ = CEF + AEF, whilst in presence of DBIMB, Cyt *b_6_f*-dependent CEF is inactivated, with TEF_DCMU+DBIMB_ = AEF.

### N_2_ fixation activity

To evaluate nitrogenase activity, an acetylene reduction assay was conducted in anoxic conditions. 2 mL of 4 days-old cultures of *Anabaena*, grown in different environmental conditions, were resuspended at OD_750_ 0.3 and placed in 8 mL gas chromatography glass vials in air + 10% acetylene (v/v). Vials were incubated for 3.5 hours upon the same environmental conditions of the original cultures before the quantification of acetylene to ethylene conversion through gas chromatography. Experiments were performed on an Agilent Technologies 8860 GC system coupled with flame ionization detector (FID) and a thermal conductivity (TCD) detector. The system was equipped with a HP-PLOT U and a MS-5A column, Argon carrier gas flow was 6 mL min^-1^ at a constant pressure of 11.121 psi, the FID and TCD detectors were kept at 250 °C. Oven was kept at the constant temperature of 40 °C for 6.5 minutes. Headspace samples were injected using PAL3 series 2 Autosampler Systems equipped with a gas-tight syringe (SGE autosampler syringe) injecting 100 μL. Experiments were performed at least in triplicate. No over-hydrogenation to ethane was detected based on the detection limit of our instrument. Calibration curves, performed in triplicate, were collected by injecting known quantities of gas mixture standards containing either ethylene (99.95 vol.% or 2.5 vol.% or 30 vol.%) or acetylene (2.5 vol.% or 1 vol.%) (Supplementary Fig. S3a and S3b). An example of a typical gas chromatogram for the reduction reaction of acetylene to ethylene is reported in Supplementary Figure S3c. Nitrogenase activity was expressed as nmol of ethylene produced per unit of time (h^-1^) and Chl content (µg Chl^-1^).

### Statistical Analysis

To describe the statistical significance of the impact of the different experimental variables (i.e. light, CO_2_ and N availability) on the quantitative measures of the different physiological parameters (e.g. duplication rate, TEF, P700 and pigment content) a conditional inference-based recursive partitioning tree, implemented in the R package “party” (Hothorn *et al*., 2006), was performed.

In detail, partitioning tree is generally used for a graphical representation of the interaction of different predictor’s variables. The approach exploits the experimental variable with the lowest *p*-value (generally significant after Bonferroni correction) as the first node of the decision tree. Then, two subgroups are generated, both characterized by different values of experimental determinations of physiological parameters. For both subgroups, the experimental variable with the lowest significant p-value is taken as the second or third node. The final tree model shows the splitting in each node, according to the experimental variables with the highest statistical significance.

Moreover, the intensity of correlation between the quantitative responses of the physiological parameters was described in all the different environmental conditions tested. First, the Pearson’s correlation coefficient was estimated and then correlograms using the R package “corrplot” were built (Friendly, 2002).

## Results

### Growth and N_2_ fixation respond differently to light and CO_2_ availability

*Anabaena* sp. PCC 7120 (hereafter *Anabaena*) was grown in different N, light and CO_2_ conditions. Light and CO_2_ are the metabolic inputs driving photosynthetic C assimilation, while the N availability in the medium (e.g. NO_3_^-^) controls the differentiation of VCs into HCs and therefore the establishment of the N_2_ fixation ability.

The response of the biological system to the different metabolic inputs was firstly assessed monitoring growth (OD_750_) (Fig. 1a and S4a). In all conditions tested the growth rate was stimulated by excess CO_2_, regardless of irradiance and the N availability (Fig. 1b), making the CO_2_ availability the environmental parameter with the predominant effect on growth (Supplementary Figure S4b) and showing that in the conditions tested the growth rate was dependent on the photosynthetic activity. In diazotrophic conditions cells always showed a slower growth, compared with similar conditions upon NO_3_^-^ repletion, suggesting a decrease in bioavailable N to the central metabolism. In atmospheric CO_2_, light intensity did not show a significant effect, suggesting cells were not able to fully exploit the available radiation because of a CO_2_ limitation (Fig. 1b). Here, N availability had a minimal effect on growth, suggesting that cells were fully able to fulfil the N demand also by N_2_ fixation.

**Fig. 1:**
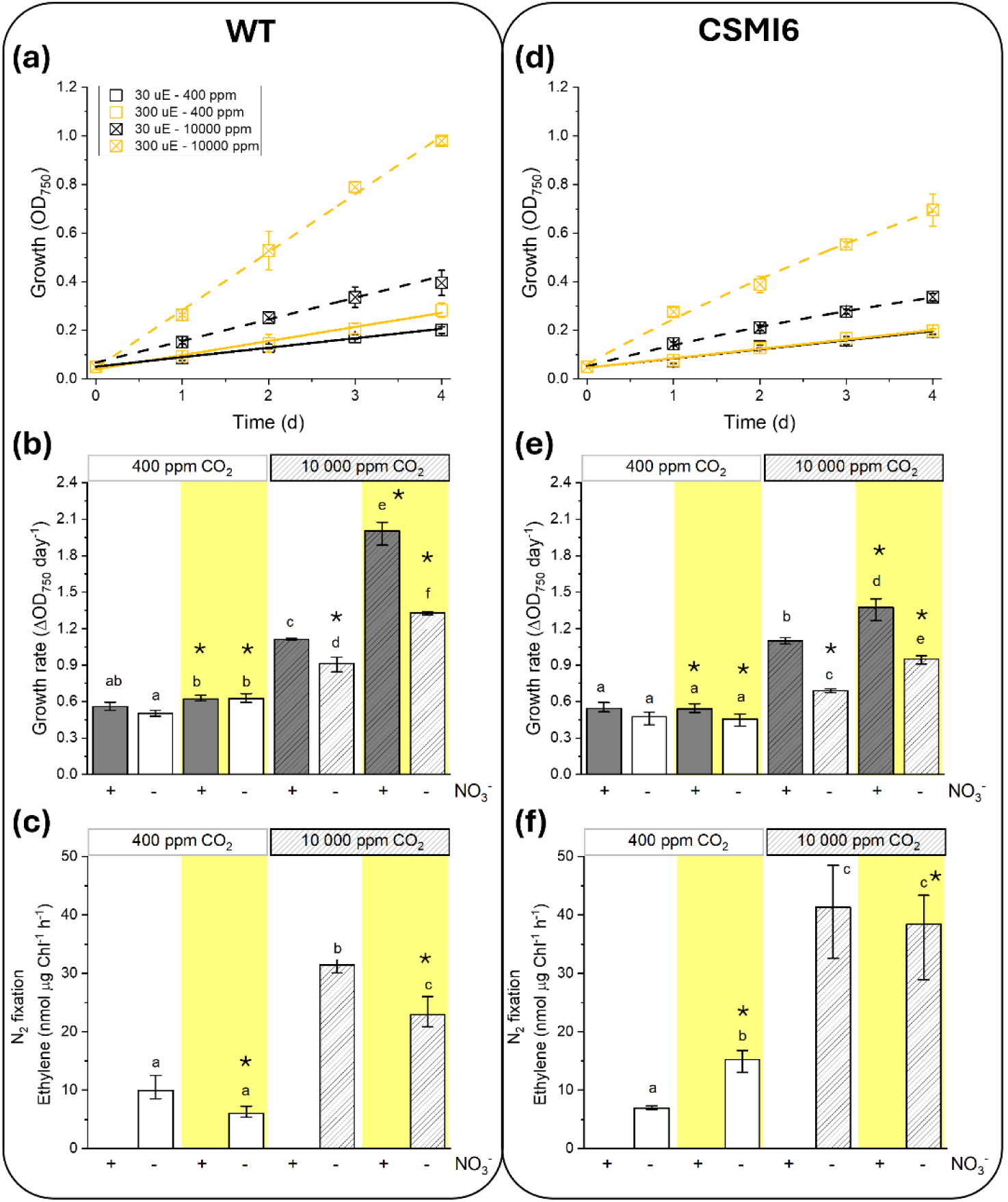
Growth and N_2_ fixation of *Anabaena* upon different light and CO_2_ conditions. (a) Growth, expressed as OD_750_, in diazotrophic conditions upon different CO_2_ and light availability for *Anabaena* WT. Growth curves upon NO_3_^-^ replete conditions are shown in Supplementary Figure S4a. (b) Growth rates of *Anabaena* WT, calculated from data in Fig. 1a and Supplementary Fig. S4a. (c) N_2_ fixation activity, expressed as nmol of ethylene converted per unit of time (h^-1^) and Chlorophyll content (µg Chl^-1^), in different N, CO_2_ and light regimes for *Anabaena* WT. (d) Growth in diazotrophic condition (growth curves upon NO_3_^-^ replete conditions shown in Supplementary Figure S4d), (e) growth rates, calculated from data in Fig. 1d and Supplementary Figure S4d and (f) N_2_ fixation activity of *Anabaena* mutant CSMI6. Grey and empty bars represent samples in NO_3_^-^ replete and diazotrophic conditions, respectively. White and yellow backgrounds indicate samples grown upon low (30 µmol photons m^-2^ s^-1^) or high (300 µmol photons m^-2^ s^-1^) irradiance, respectively; full colours and fill pattern indicate samples grown in atmospheric (400 ppm) or excess (10000 ppm) CO_2_, respectively. Data represents averages of > three biological replicates (± SD). Different lowercase letters and asterisks indicate statistically significant differences between samples upon different environmental conditions within the same strain and between strains upon the same environmental condition, respectively (one-way ANOVA, p < 0.05).

The N_2_ fixation activity in all cells was quantified through the reduction of acetylene to ethylene *via* gas chromatography (Supplementary Fig. S4c). Overall, the N_2_ fixation activity, like growth, was higher in high CO_2_. However, the correlation with growth was only partial, as the higher light did not induce a higher N_2_ fixation capacity. The greatest activity was measured upon low light and excess CO_2_, and it dropped with stronger irradiance. Also in atmospheric CO_2_, upon low irradiance the N_2_ fixation activity was higher than in high light (Fig. 1c). It is worth noting that the quantification of the acetylene reduction was performed in atmospheric CO_2_. Here cultures could have experienced CO_2_ limitation, overall curbing photosynthesis and underestimating the N_2_ fixation activity (Fig. 1c), especially upon high irradiance where also photoinhibition occurs.

To investigate further the connection between growth and N_2_ fixation, an *Anabaena* mutant strain, i.e. CSMI6, impaired in the vehiculation of fixed N to VCs (Burnat *et al*., 2014), was exposed to the same experimental conditions and both growth (Fig. 1d) and N_2_ fixation were compared to the wild-type strain (wild-type, WT).

In strain CSMI6, growth followed the same trend of the WT, increasing with the CO_2_ availability, regardless of irradiance, in both N regimes (Fig. 1e). However, the mutation drove a significant growth reduction with respect to the WT strain (Fig. 1d and Supplementary Fig. S4d), with cells upon high irradiance and excess CO_2_ experiencing the strongest decrease (i.e. ∼30 %) (Fig. 1e). The mutation therefore impaired the cells growth, likely by affecting the ability to exploit the N fixed by HCs. The N_2_ fixation also showed the greatest values upon excess CO_2_, like in the WT strain. The mutant showed a general increase of the N_2_ fixation capacity with respect to the WT in most of the conditions tested, confirming that the phenotype was not due to the alteration of N_2_ fixation, rather to the transport of the molecules (Fig. 1f).

### Impact of N_2_ fixation on the composition and activity of the photosynthetic machinery

To investigate the role of the photosynthetic machinery on the N_2_ fixation of *Anabaena*, both its composition and functionality were assessed in the conditions tested above.

In the WT strain the chlorophyll *a* (Chl) content overall significantly decreased upon diazotrophic with respect to NO_3_^-^ replete conditions (Fig. 2a). The chlorophyll/carotenoid (Chl/Car) ratio instead remained constant (Supplementary Fig. S5a), suggesting that cells generally accumulate less pigment binding proteins when the N_2_ fixation is active. Moreover, upon diazotrophic conditions the Chl content significantly decreased with irradiance, regardless of the CO_2_ availability (Fig. 2a). High irradiance induced a reduction of the Chl/Car ratio regardless of the N regime (Supplementary Fig. S5a), thus suggesting a relative increase of the Car content, a common response to excess illumination.

**Fig. 2:**
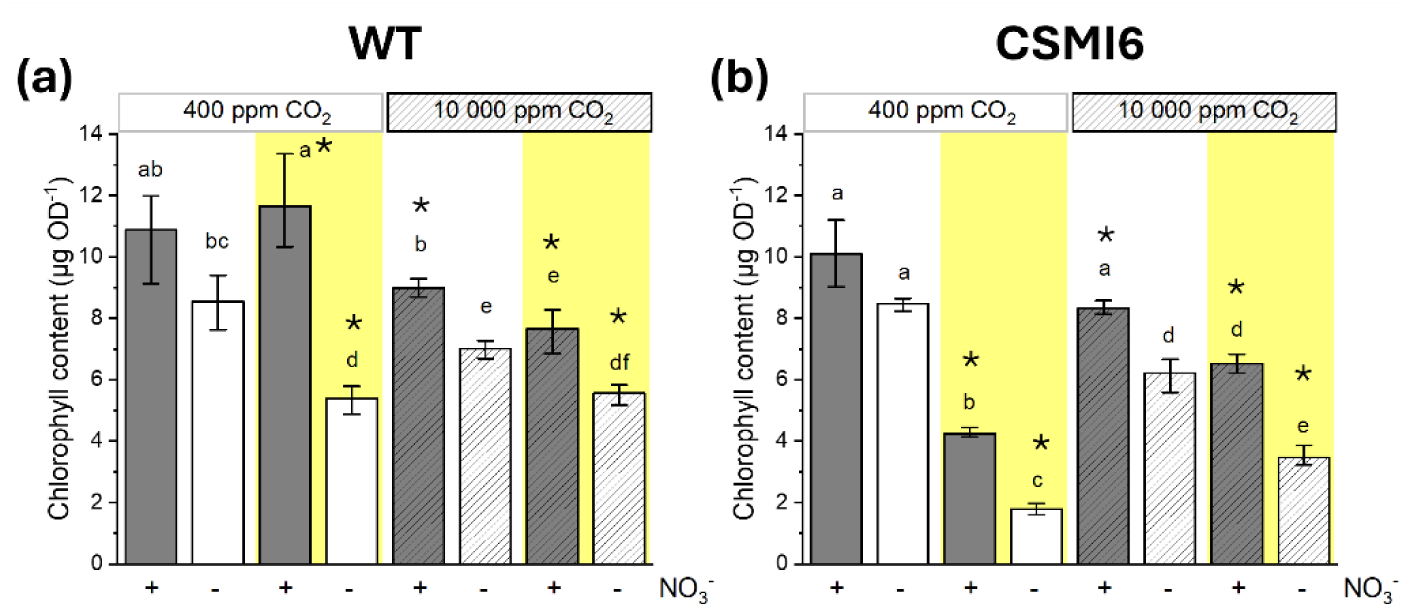
Chlorophyll content of *Anabaena* upon different irradiances. Chlorophyll content per unit of biomass (OD^-1^) upon different irradiances and CO_2_ availability for *Anabaena* WT (a) and mutant CSMI6 (b), comparing NO_3_^-^ replete with diazotrophic conditions. Grey and empty bars indicate NO_3_^-^ replete and diazotrophic conditions, respectively; white and yellow backgrounds indicate limiting (30 µmol photons m^-2^ s^-1^) or excess (300 µmol photons m^-2^ s^-1^) irradiance, respectively; full colours and fill pattern indicate limiting (400 ppm) or excess (10000 ppm) CO_2_, respectively. Data represents averages of at least three biological replicates (± SD). Different lowercase letters and asterisks indicate statistically significant differences between samples upon different environmental conditions within the same strain and between strains upon the same environmental condition, respectively (one-way ANOVA, p < 0.05).

In the mutant strain, the Chl content overall decreased with irradiance more than in the WT strain (∼50% vs < 25%, respectively) (Fig. 2b). Interestingly, the Chl/Car ratio also significantly decreased (Supplementary Fig. S5b), indicating that the Car content too was affected by the mutation, as confirmed by the analysis of variance (Table S1).

The composition of the photosynthetic apparatus was further assessed exploiting the JTS-10 absorbance spectroscopy, measuring the P700 absorbance signal at 705 nm. P700 is a molecular dimer of chlorophyll *a* associated with the reaction-centre of photosystem I (PSI) and it is the PSI primary electron donor. Upon dark exposure, photosystem I (PSI) is fully reduced (Supplementary Fig. S1a and Fig. 3a). The switch on of an orange light drives to its oxidation (P700^+^) (Fig. 3a: top orange bar) and to a consequent change of absorption of the red fraction of the light spectrum (Supplementary Fig. S1a grey straight line). In presence of two PSII and cytochrome *b_6_f* (Cyt *b_6_f*) inhibitors, 3-(3,4-dichlorophenyl)-1,1-dimethylurea (DCMU) and 2,5-Dibromo-6-isopropyl-3-methyl-1,4-benzoquinone (DBIMB), respectively (Supplementary Figs. S1a and S2), the electron flux towards PSI is blocked, thus enabling to reach the maximal oxidation of the PSI reaction centres upon light exposure. The maximal P700^+^ signal obtained in these conditions (P700_max_) depends on the amount of PSI reaction centres present in the sample and can be exploited for the quantification of the content of active PSI units.

**Fig. 3:**
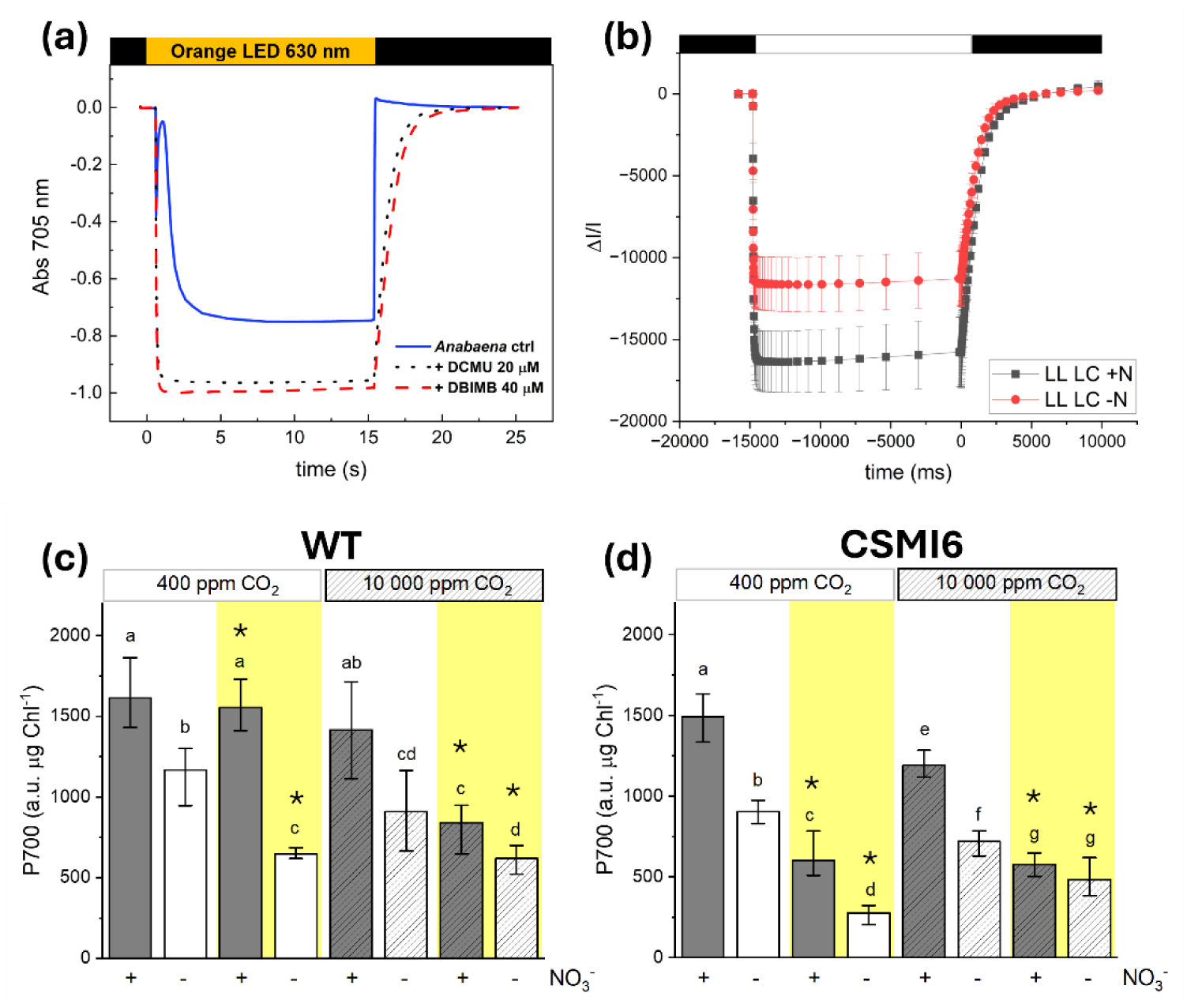
P700 absorption spectroscopy and content of active PSI units in *Anabaena*. (a) Representative traces of P700 absorption (705 nm) of *Anabaena* WT (blue continuous line) upon exposure for 15 s to saturating light (orange top bar) in presence of the PSII inhibitor DCMU (20 µM, black dot line) and Cytochrome *b_6_f* inhibitor DBIMB (40 µM, red dashed line). (b) Representative traces for the PSI quantification for WT cells upon low light and atmospheric CO_2_, comparing NO_3_^-^ replete (black) with diazotrophic (red) conditions. The PSI content was evaluated from the maximum absorption of P700^+^ at 705 nm, in the presence of DCMU and DBMIB and upon a saturating light of 2080 µmol photons m^-2^ s^-1^ (see methods for details). White box, actinic light on; black box, actinic light off. Representative traces for the other environmental conditions tested in this work are shown in Supplementary Figure S5. (c-d) Maximum P700 oxidized measured upon exposure to different irradiances and CO_2_ availability for *Anabaena* WT (c) and CSMI6 (d), comparing NO_3_^-^ replete with diazotrophic conditions. Grey and empty bars indicate NO_3_^-^ replete and diazotrophic conditions, respectively; white and yellow backgrounds indicate limiting (30 µmol photons m^-2^ s^-1^) or excess (300 µmol photons m^-2^ s^-1^) irradiance, respectively; full colours and fill pattern indicate limiting (400 ppm) or excess (10000 ppm) CO_2_, respectively. Data represents averages of at least three biological replicates (± SD). Different lowercase letters and asterisks indicate statistically significant differences between samples upon different environmental conditions within the same strain and between strains upon the same environmental condition, respectively (one-way ANOVA, p < 0.05).

This method has been applied to plants and eukaryotic microalgae, with only few examples in cyanobacteria (Bailey *et al*., 2008; Liberton *et al*., 2022), but to the best of our knowledge never in *Anabaena*. The P700_max_ signal was observed to be linearly dependent on the cell concentration (Supplementary Fig. S1b). Moreover, the fraction of oxidised P700 (P700^+^/P700) was proportional to the irradiance and their relationship could be described by a 2-variable exponential function, with the number of active PSIs increasing with irradiance upon light limitation, whilst reaching saturation at higher irradiances (Supplementary Fig. S1c), highlighting the P700_max_ absorption was indeed a reliable probe to infer quantitative information on the content of PSI units in the host.

Cells of the WT strain showed overall a lower PSI content in diazotrophic than in NO_3_^-^ replete conditions (Fig. 3b and Supplementary Fig. S6), with ∼50% less active PSI units in diazotrophy (Fig. 3C). Moreover, upon diazotrophic conditions the content of the active PSI units significantly decreased with irradiance, similarly to the Chl content (Fig. 2a). The mutant strain overall showed a lower PSI content with respect to the WT in all tested conditions (Supplementary Fig. S6), with the largest difference upon high light (Fig. 3d), confirming the negative effect of the mutation on the accumulation of photosynthetic proteins.

The P700 absorption signal can also be exploited to obtain information on the PSI reduction and oxidation rates and the rate of P700 re-reduction when light is switched off can be used to quantify the Total Electron Flow (TEF) (Fig. 3a and 4a) (see methods for details). The reliability of this approach in *Anabaena* was first assessed by verifying that the P700^+^ reduction rate in the dark was proportional to the irradiance and their relationship could be described by a 2-variable exponential function (Supplementary Fig. S1d). The total photosynthetic activity (TEF) of the biological system linearly increased with irradiance upon light limitation, whilst at higher values it reached saturation (Supplementary Fig. S1e), as expected (Sun & Wang, 2018), highlighting that the P700 absorption is a robust probe to infer quantitative information on the photosynthetic functionality of the host.

**Fig. 4:**
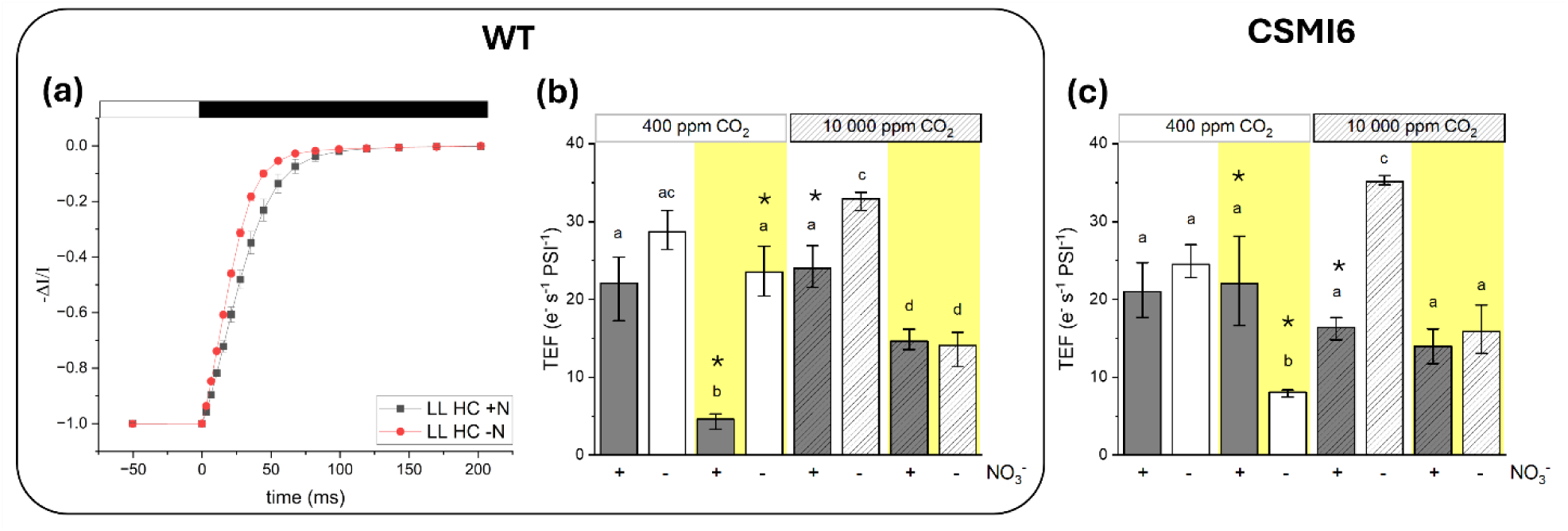
Total Electron Flow in *Anabaena*. (a) Representative traces of oxidized P700 re-reduction kinetics for WT cells upon low light and excess CO_2_, comparing NO_3_^-^ replete (black) with diazotrophic (red) conditions. The kinetics were used for the total electron transport (TEF) quantification (see methods for details) upon exposure to a saturating light intensity (2080 µmol photons m^-2^ s^-1^) before evaluating the recovery kinetics in the dark, here reported. White box, actinic light on; black box, actinic light off. Representative traces for the other environmental conditions tested in this work are shown in Supplementary Figures S6 and S7. (b-c) Total electron flow (TEF) per unit of PSI and time (s^-1^) upon exposure to different irradiances and CO_2_ availability for *Anabaena* WT (b) and *Anabaena* mutant CSMI6 (c), comparing NO_3_^-^ replete with diazotrophic conditions. Grey and empty bars indicate NO_3_^-^ replete and diazotrophic conditions, respectively; white and yellow backgrounds indicate limiting (30 µmol photons m^-2^ s^-1^) or excess (300 µmol photons m^-2^ s^-1^) irradiance, respectively; full colours and fill pattern indicate limiting (400 ppm) or excess (10000 ppm) CO_2_, respectively. Data represents averages of at least three biological replicates (± SD). Different lowercase letters and asterisks indicate statistically significant differences between samples upon different environmental conditions within the same strain and between strains upon the same environmental condition, respectively (one-way ANOVA, p < 0.05).

In most cases, cells of the WT strain showed overall faster re-reduction kinetics of P700^+^ in diazotrophy with respect to NO_3_^-^ replete conditions (Fig. 4a and Supplementary Fig. S7). The total photosynthetic activity showed a significant increase in diazotrophy with respect to NO_3_^-^ repletion, suggesting a higher electron transport activity (Fig. 4b). The only exception to this trend were cells exposed to light and CO_2_ excess, where TEF was lower and similar in both N regimes (Fig. 4B), in contrast to the growth data that showed instead the greatest values measured in this work (Fig. 1b). This is likely dependent on the fact that the JTS-10 absorbance spectroscopy employed in this work was performed upon atmospheric CO_2_, thus likely underestimating the total photosynthetic capacity in this condition.

Interestingly, upon NO_3_^-^ repletion TEF significantly decreased with irradiance, regardless of the CO_2_ availability, with the strongest reduction upon high light and atmospheric CO_2_ (Fig. 4b). In contrast, upon diazotrophic conditions TEF significantly decreased with irradiance only upon excess CO_2_, whilst maintained stable values upon atmospheric CO_2_ conditions (Fig. 4b), suggesting that the establishment of the N_2_ fixation capacity helps avoiding the saturation of the photosynthetic electron transport chain.

Surprisingly, the effect of the mutation in the strain CSMI6 on TEF was only observed upon high light and atmospheric CO_2_, in which, differently from the WT, it showed a high activity upon NO_3_^-^ repletion whilst significantly decreasing in diazotrophic conditions (Fig. 4C).

### Impact of N_2_ fixation on different photosynthetic electron pathways

TEF depends on the contribution of three components: i) a PSII-dependent linear electron flow (LEF), and two PSII-independent electron pathways, namely ii) cyclic (CEF) around PSI and iii) other alternative flows (AEF), feeding or taking electrons from the photosynthetic electrons transport chain. The contribution of these three components to the total photosynthetic activity can be estimated using specific inhibitors. When cells are treated with DCMU, PSI oxidation levels and reduction rates are determined only by PSII-independent processes, such as CEF and AEF. In the presence of both DCMU and DBIMB, instead, only AEF determines the total photosynthetic activity (Fig. 3a and Fig. 5a). To investigate whether the increased photosynthetic capacity upon N_2_ fixation was the result of an altered contribution of specific photosynthetic electron pathways, the PSII-dependent (i.e. LEF) and PSII-independent (i.e. CEF and AEF) contributions were assessed.

**Fig. 5:**
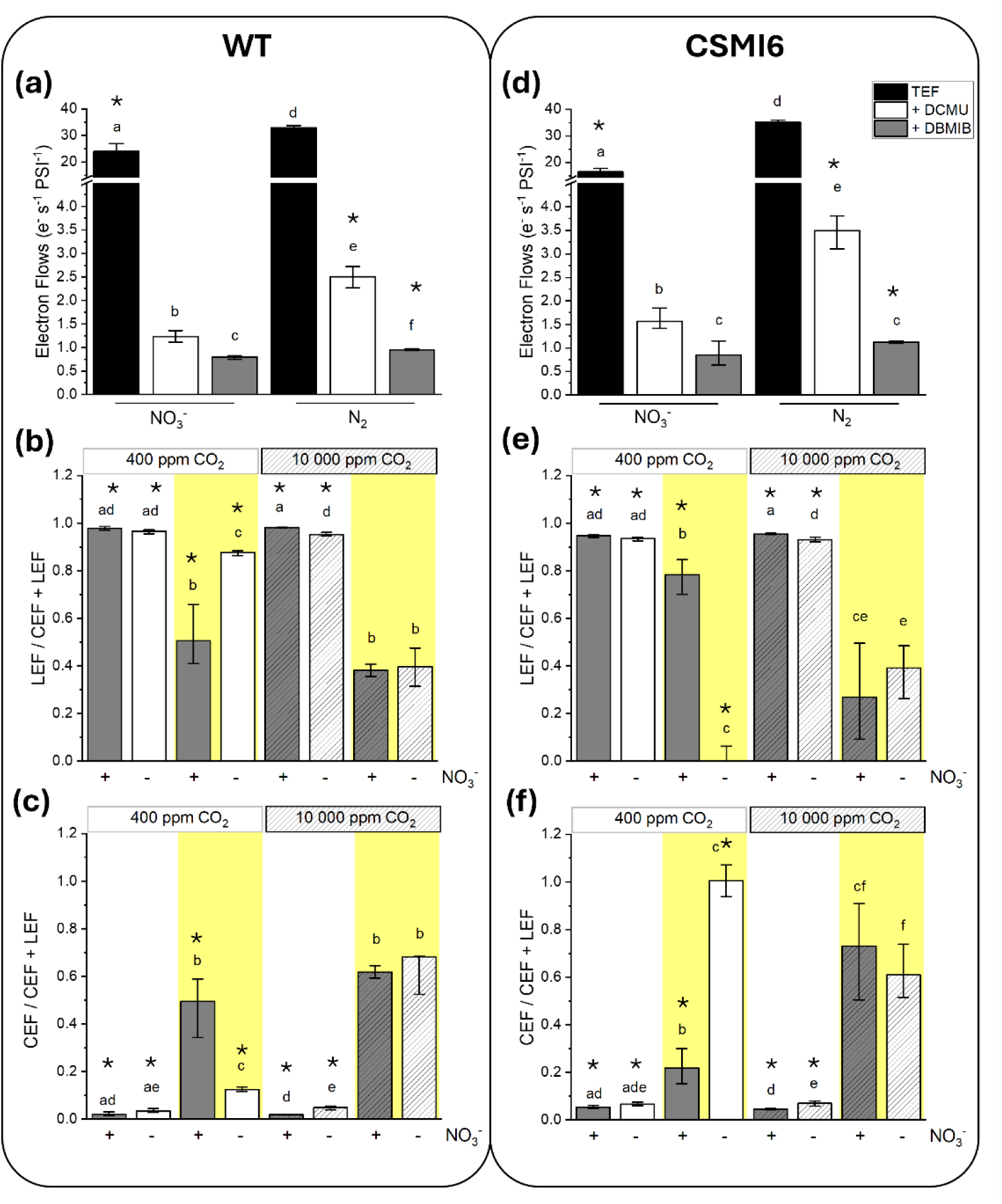
Quantification of the photosynthetic electron flows in *Anabaena*. (a) Quantification of P700^+^ re-reduction rates after illumination in untreated (black), DCMU-treated (white) and DCMU and DBMIB-treated samples (grey) for *Anabaena* WT. *Anabaena* cells upon low light and excess CO_2_, comparing NO_3_^-^ replete with diazotrophic conditions. Black bar, total electron flow; White bar, contribution of cyclic and other alternative electron flows; Grey bar, other alternative electron flows. The subtraction of the white from the black bar shows the linear electron flow, while the grey from the white bar shows the cyclic electron flow and such calculations are reported in the other panels of the figure. Quantification of linear (LEF) (b) and cyclic (CEF) (c) electron flows per unit of PSI and time (s^-1^) upon exposure to different irradiances and CO_2_ availability for *Anabaena* WT, comparing NO_3_^-^ replete with diazotrophic conditions. (d) Quantification of P700^+^ re-reduction rates for *Anabaena* mutant CSMI6. Quantification of (LEF) (e) and CEF (f) per unit of PSI and time upon exposure to different irradiances and CO_2_ availability for *Anabaena* CSMI6. Grey and empty bars indicate NO_3_^-^ replete and diazotrophic conditions, respectively; white and yellow backgrounds indicate limiting (30 µmol photons m^-2^ s^-1^) or excess (300 µmol photons m^-2^ s^-1^) irradiance, respectively; full colours and fill pattern indicate limiting (400 ppm) or excess (10000 ppm) CO_2_, respectively. Data represents averages of at least three biological replicates (± SD). Different lowercase letters and asterisks indicate statistically significant differences between samples upon different environmental conditions within the same strain and between strains upon the same environmental condition, respectively (one-way ANOVA, p < 0.05).

The PSII-dependent photosynthetic activity was the major contributor to TEF in both N regimes (Fig. 5a). Upon treatment with DCMU we observed that the contribution of CEF was instead only marginal with respect to LEF in NO_3_^-^ repletion, while it significantly increased upon N_2_ fixation. On the other hand, AEF showed overall values < 1 e^-^ s^-1^ PSI^-1^, regardless of the N regime, making its contribution on the total photosynthetic functionality of *Anabaena* negligible in all the conditions tested (Fig. 5a). Interestingly, the efficiency of LEF and CEF did not change significantly upon N_2_ fixation with respect to NO_3_^-^ repletion, with the only exception upon atmospheric CO_2_ and high light (Fig. 5b and 5c). Here LEF significantly decreased in high light upon NO_3_^-^ repletion (Fig. 5b) and the reduction was compensated by a strong increase of the CEF efficiency (Fig. 5c), likely to activate photoprotection from high light. On the contrary, upon diazotrophic conditions LEF retained its efficiency (Fig. 5b) and CEF was not significantly activated (Fig. 5c), suggesting that the establishment of the N_2_ fixation capacity likely enables to dissipate enough excitation energy. On the other hand, in excess CO_2_ the efficiency of CEF exceeded that of LEF upon high light, in both N regimes (Fig. 5c), suggesting that PSII-independent photosynthetic electron pathways became predominant.

In the mutant strain CSMI6, LEF was overall only marginally affected by the mutation and remained the major contribution to TEF in both N regimes. Interestingly, CEF significantly increased upon N_2_ fixation with respect to the WT (Fig. 5d), suggesting a major effect of the mutation on the PSII-independent photosynthetic activity when photosynthesis segregation occurs, as also confirmed by the analysis of variance (Table S1). This effect was maximal upon high light and atmospheric CO_2_, in which LEF was suppressed and CEF instead reached the maximal efficiency (Fig. 5e and 5f).

### Correlation analysis between photosynthetic parameters

The data collected so far showed that the contribution of specific electron pathways on the total photosynthetic activity in diazotrophy changed the most upon high light exposure, regardless of the CO_2_ availability (Fig. 5). To clarify the implications of the observed modifications on the physiology of *Anabaena*, a correlation analysis was run upon high light exposure in N_2_ fixing conditions (Fig. 6a).

**Fig. 6:**
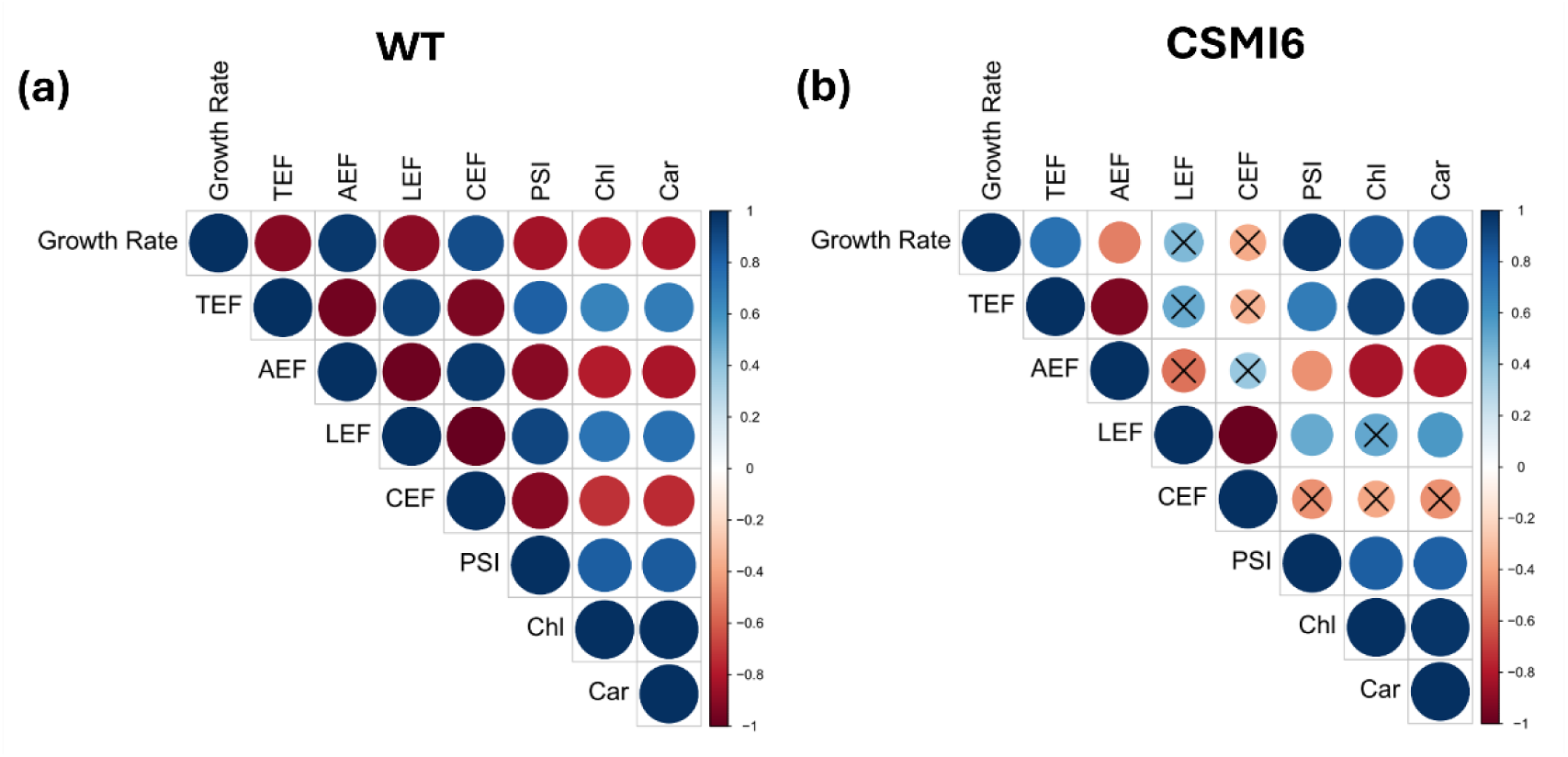
Correlation analysis between physiological parameters. (a-b) Correlograms showing correlation values between different physiological parameters upon excess light in diazotrophic conditions, regardless of the CO_2_ availability, in *Anabaena* WT (a) and mutant CSMI6 (b). Blu, positive correlation; Red, negative correlation. Crosses indicate that the correlation value is not statistically significant. The numbers on the colour scale on the right of each panel indicate the level of Pearson’s correlation. TEF, Total Electron Flow; LEF, Linear Electron Flow; CEF, Cyclic Electron Flow; AEF, Alternative Electron Flow; P700, PSI content; Chl, Chl content; Car, Car content.

In the WT strain, TEF positively correlated with the content of both pigments and active PSI units, indicating its dependence on the components of the photosynthetic machinery, as expected. TEF also positively correlated with LEF, suggesting that the total photosynthetic functionality is overall mainly determined by the PSII-dependent activity (Fig. 6a). Surprisingly the growth rate negatively correlated with TEF, LEF and the content of pigments and active PSI units, whilst it positively correlated with both CEF and AEF (Fig. 6a), highlighting that also the PSII-independent photosynthetic activity can play a role in the energetic balance of the organism.

In the mutant strain we did not observe major differences in the correlogram between physiological parameters. The only exception was for the growth rate that inverted its correlation trend with respect to the WT and showed a positive correlation with TEF, LEF and the content of the components of the photosynthetic machinery indeed, whilst it showed a negative correlation with both CEF and AEF (Fig. 6b), overall confirming that the impact of the mutation on growth involves a major remodelling of the photosynthetic functionality.

## Discussion

### The rate of provision of fixed N to photosynthesis can limit growth

The experimental set-up of this work enabled to characterize *Anabaena* cultures in different environmental conditions. Upon NO_3_^-^ repletion, the growth of *Anabaena* increased with the availability of both light and CO_2_ (Fig. 1b), suggesting its dependence on the photosynthetic C assimilation, as expected in strict photoautotrophic conditions (Malatinszky *et al*., 2017). Upon atmospheric CO_2_ growth was relatively slow and did not increase with irradiance (Fig. 1b), suggesting that photosynthesis was limited by the C availability (Table 1). The exposure to higher light with limiting CO_2_ thus means that the cells were experiencing a strong light excess (Table 1), as they were not able to use the absorbed energy for growth. Under high CO_2_ supply, the growth was clearly faster and it was also able to take advantage from stronger irradiance (Fig. 1b), suggesting instead a limitation by the availability of the light energy (Table 1). As expected, *Anabaena* cultures were still able to grow in the absence of a combined N source in the medium (Fig. 1a), consistent with its ability to develop HCs and fix atmospheric N_2_ (Berman-Frank *et al*., 2001; Herrero *et al*., 2016; Forchhammer & Selim, 2019; Nieves-Morión *et al*., 2021). In diazotrophic conditions, the growth of *Anabaena* increased following the availability of both light and CO_2_ (Fig. 1b), indicating its major dependence on the photosynthetic C assimilation also when the N_2_ fixation was active. Also, the N_2_ fixation activity increased upon excess CO_2_ (Fig. 1c), therefore suggesting it depends on the photosynthetic C assimilation as well.

**Table 1:**
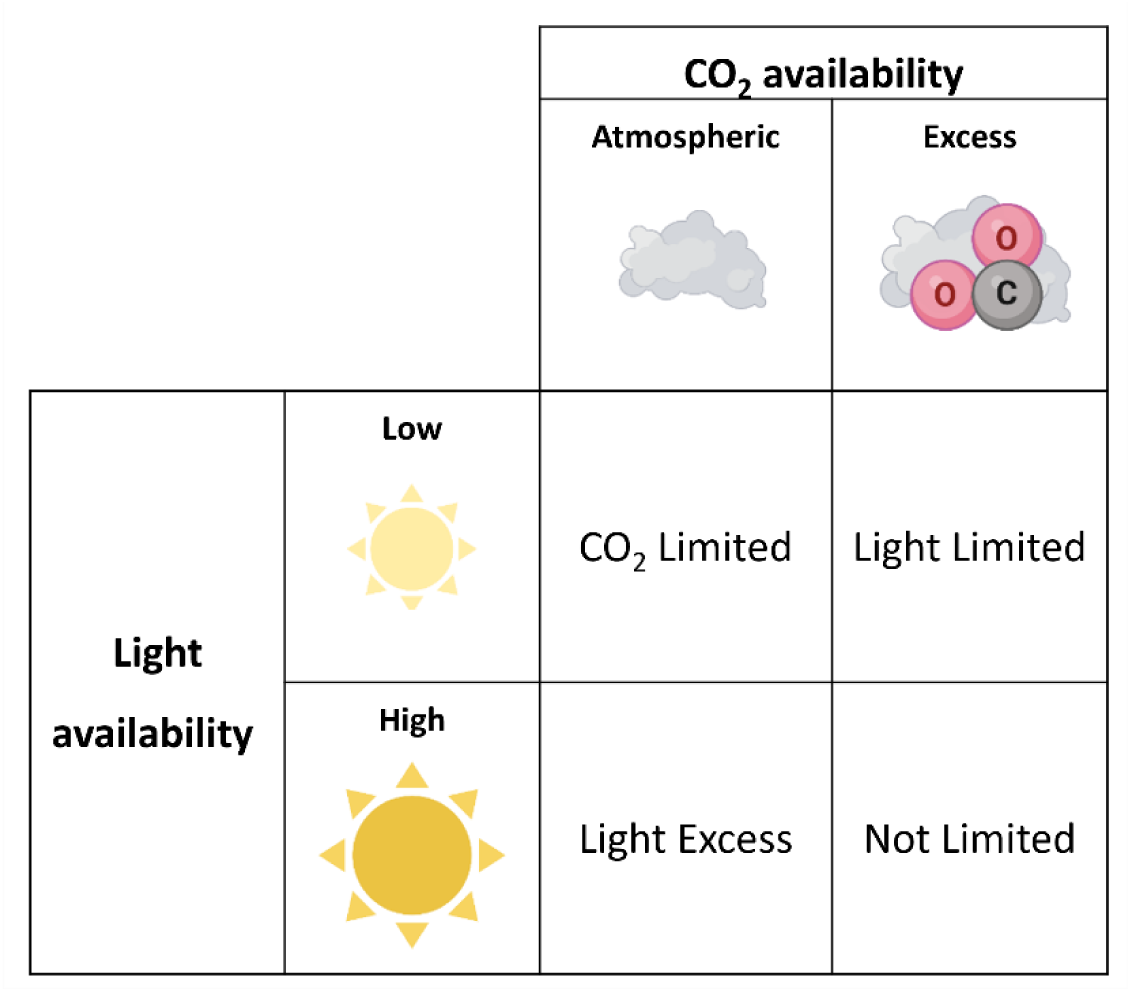
Classification of photosynthesis in the four environmental conditions of this work. Atmospheric and excess CO_2_ corresponds to 400 ppm and 10000 ppm, respectively. Low and high light corresponds to 30 and 300 µmol photons m^-2^ s^-1^, respectively.

|  |  | CO<sub>2</sub> availability |  |
| --- | --- | --- | --- |
|  |  | Atmospheric | Excess |
| Light availability | Low | CO<sub>2</sub> Limited | Light Limited |
|  | High | Light Excess | Not Limited |

However, in excess CO_2_ cultures in diazotrophic conditions showed a significant reduction of the growth rate with respect to NO_3_^-^ repletion, suggesting that other parameters affect growth when the N_2_ fixation is active (Fig. 1b). To investigate the relationship between N_2_ fixation and growth, a mutant strain of *Anabaena* (CSMI6) with an impaired capacity of synthetising cyanophycin, a N-rich polymer of aspartate and arginine that stands at the junction between VCs and HCs and regulates the provision of fixed N to the former (Burnat *et al*., 2014; Nieves-Morión *et al*., 2021), was exploited. Upon diazotrophic conditions, cyanophycin is synthesised by cyanophycinase producing β-aspartyl arginine, which is transferred to and hydrolyzed by isoaspartyl dipeptidase in VCs, in order to make N bioavailable for the central metabolism (Burnat *et al*., 2014). CSMI6 is a knock-out mutant of the gene *all3922*, encoding for the hydrolysing enzyme, which shows an increased accumulation of cyanophycin in the form of granules. Consequently, the provision of fixed N to VCs is curbed in the mutant (Burnat *et al*., 2014).

Upon CO_2_ limitation (Table 1) the mutant strain did not show an altered growth phenotype with respect to the WT (Fig. 1e), confirming that photosynthesis was C-limited indeed and that the need for fixed N was therefore reduced, suggesting that here complementary vehiculation pathways of fixed N (e.g. glutamine and arginine) (Wolk *et al*., 1974; Thomas *et al*., 1977; Burnat *et al*., 2014) are sufficient to compensate for the absence of the flux from cyanophycin. However, in excess CO_2_ the mutant strain showed a significant reduction of growth with respect to the WT (Fig. 1e), with the largest growth difference (i.e. >30%) observed in high light, indicating an increased demand for fixed N that makes the rate of provision *via* cyanophycin essential.

Because the largest impact of the mutation on growth was observed when photosynthesis is not limited by its two main metabolic inputs (Table 1), we suggest that the central metabolism depends on the provision of fixed N from HCs but that when the speed of photosynthesis exceeds the rate of provision, the latter becomes limiting.

### The acclimation response to high light differs in the two N regimes

In nature, weather conditions and water movements drive fluctuations in the availability of light energy, generating sudden peaks of irradiance that can saturate photosynthesis and last indefinitely. Photosynthetic organisms evolved regulatory mechanisms to withstand excess irradiance, which rely on the synthesis of pigments and proteins and the repair of photosynthetic components to minimize the consequences of photodamage, whose efficiency therefore depends on the availability of N. Upon NO_3_^-^ repletion, *Anabaena* acclimated photosynthesis to high light increasing the Car content (Fig. 2a), a common response among photosynthetic organisms (Izuhara *et al*., 2020; Pagels *et al*., 2021).

In plants (Joliot & Johnson, 2011; Shikanai, 2014) and algae (Peltier *et al*., 2010; Meneghesso *et al*., 2016) the acclimation to high light involves also the remodulation of the contribution of different photosynthetic electron fluxes on the total photosynthetic activity. Pseudo-cyclic and cyclic electron pathways around PSI are induced to alleviate the overreduction of the photosynthetic electron transport chain, whereas the efficiency of the linear electron flow from PSII is reduced. A similar response is expected also in cyanobacteria as they carry the molecular complexes mediating the activation of the same mechanisms (Mullineaux, 2014). Recent works highlighted that the contribution of the cyclic electron pathway on the total photosynthetic activity can exceed 35% in cyanobacteria (Miller *et al*., 2021; Theune *et al*., 2021), overall suggesting that the remodulation of photosynthetic electron fluxes likely plays a more relevant role in cyanobacteria than plants and eukaryotic algae. However, the understanding of the contribution of different photosynthetic electron pathways on the acclimation to high light in cyanobacteria has lagged behind its expected relevance in nature, mainly because of technical limitations.

The quantification of photosynthesis in plants is commonly performed through the monitoring of the chlorophyll fluorescence signal *in vivo* (Acuña *et al*., 2016; Ogawa *et al*., 2018). In cyanobacteria the interpretation of Chl fluorescence yield data is complicated by the presence of both phycobiliproteins and respiration acting on the same membrane systems, which often prevent the exploitation of the technique for a reliable quantification of photosynthesis (Ogawa *et al*., 2018; Viola *et al*., 2019). Moreover, whilst the Chl fluorescence is a good proxy for the activity of PSII, the signal from PSI does not make a significant contribution below 700 nm, preventing the inclusion of its contribution on the total photosynthetic activity (Murchie & Lawson, 2013). To overcome these limitations, this work exploited the P700^+^ absorbance signal (Supplementary Figure S1a), a valuable probe to assess the contribution of all the electron fluxes on the photosynthetic functionality of the host, which finds only few application examples in cyanobacteria and has not been systematically applied to filamentous diazotrophic species to date (Joliot *et al*., 2004; Simionato *et al*., 2013; Storti *et al*., 2019; Viola *et al*., 2019; Liberton *et al*., 2022).

Upon NO_3_^-^ repletion *Anabaena* showed a reduction of the total photosynthetic activity with increased irradiance (Fig. 4b) and a remodulation of the contribution of specific electron fluxes (Fig. 5). The efficiency of the PSII-dependent linear electron pathway significantly decreased with irradiance (Fig. 5b). The pseudo-cyclic contribution to the total photosynthetic activity was observed to be dispensable in *Anabaena*, suggesting a marginal role in the protection from prolonged exposure to high light, as expected (Allahverdiyeva et al. (2011)). Instead, the cyclic electron flow around PSI increased, enabling the consumption of up to 60 % of harvested photons (Fig. 5c) and pointing to the activation of a regulatory mechanism to alleviate the overreduction of the photosynthetic electron transport chain (Nogales *et al*., 2012; Shaku *et al*., 2016). Overall, this work provides evidence for a conserved role of the regulation of photosynthetic electron pathways *in vivo* to respond to changing light conditions also in filamentous diazotrophic cyanobacteria.

Interestingly, the mutation in strain CSMI6 drove to significant alterations on the acclimation to high light upon NO_3_^-^ repletion, suggesting an extensive impact on the physiology of photosynthesis. Upon high light the mutant strain showed a significant reduction of the Chl and PSI content (Fig. 2b and 3c), as well as a remodulation of the efficiency of different photosynthetic electron pathways (Fig. 5), suggesting that the acclimation of photosynthesis is impaired. These data suggest the evolution of a luxury uptake mechanism in *Anabaena* that enables to store N in the form of cyanophycin granules. In the mutant strain this N pool is not accessible to the central metabolism, causing a reduced acclimation efficiency, affecting growth upon light excess conditions (Fig 1E).

Interestingly, upon N_2_ fixation conditions the acclimation of the WT *Anabaena* strain to high light was altered. Cells in fact responded with a reduction of the content of both Chl and active PSI units (Fig. 2a and 3c) leaving unaffected the Car pool (Supplementary Fig. S5a), an inverted trend with respect to NO_3_^-^ replete conditions. Because of the N content of the Chl molecule, its reduction is a reasonable functional response to limit the chance of photosynthesis saturation upon high light, whilst responding to a limitation in the bioavailability of N upon diazotrophic conditions, as discussed above.

Upon atmospheric CO_2_ photosynthesis was maintained fully active in the WT *Anabaena* strain even when irradiance increased (Fig. 4b), suggesting that the establishment of the N_2_ fixation capacity likely provides an advantage to withstand excess irradiance. Nitrogenase indeed represents another electron sink, beside the Calvin-Benson-Bassham (CBB) cycle, for the host. If the CBB cycle calls for the vehiculation of 16 electrons along the photosynthetic electron transport chain to fix one molecule of CO_2_, the nitrogenase needs up to 80 electrons to fix one molecule of N_2_ (Seefeldt *et al*., 2018; Rapson & Wood, 2022), overall significantly increasing the demand for reducing equivalents and therefore the electron sink capacity of the biological system indeed. Upon diazotrophic conditions, TEF significantly increased with respect to NO_3_^-^ replete cultures indeed (Fig. 4b). It is however worth remembering that in this work the photosynthetic activity was normalized to the content of active PSI units. The latter decreased upon diazotrophic conditions (Fig. 3c), therefore suggesting that the total photosynthetic capacity was not affected when the N_2_ fixation was active. Nevertheless, the biological system responded by speeding up photosynthesis, confirming a greater electron sink capacity. The greatest impact was measured upon atmospheric CO_2_ and high light (Fig. 4b), namely when the CBB cycle was limited by the C availability and photosynthesis was likely saturated (Table 1). Here NO_3_^-^ replete cultures showed a 4-fold reduction of TEF with respect to low light (Fig. 4b), indeed confirming photoinhibition. On the other hand, in diazotrophic cultures, the total photosynthetic functionality was not affected, suggesting that the electron sink capacity of nitrogenase can be beneficial to withstand excess light (Fig. 4b).

Therefore, upon diazotrophic conditions the nitrogenase complex can interestingly represent a further electrons relief valve beside the cyclic electron pathways around PSI to prevent the overreduction of the photosynthetic electron transport chain upon high light.

### Regulation of photosynthetic electron pathways upon diazotrophic conditions

With the only exception of high light and excess CO_2_, upon diazotrophic conditions LEF overall represented the major contributor to TEF in the WT *Anabaena* strain (Fig. 5a-c), suggesting that the PSII-dependent photosynthetic activity of VCs plays a relevant role in feeding the nitrogenase in HCs. The large electron sink capacity of nitrogenase (Seefeldt *et al*., 2018; Rapson & Wood, 2022) is overall expected to significantly increase the demand for reducing equivalents, explaining a substantial dependence on the linear photosynthetic electron flow of VCs. This was also in line with the correlation analysis that showed that TEF positively correlated with LEF and both the content of active PSI units and pigments (Fig. 6a), suggesting that the major contribution to the total photosynthetic activity comes from VCs indeed (Wolk & Simon, 1969). These data strengthen the hypothesis that carbohydrates synthesized because of the PSII-dependent photosynthesis are likely translocated from VCs and metabolized in HCs *via* the oxidative pentose pathway, generating NADPH (Bottomley & Stewart, 1977; Summers *et al*., 1995; Magnuson, 2019). The latter is then suggested to feed reducing equivalents to the proton-pumping NDH-1 complex, which in turn provides electrons to the nitrogenase (Magnuson, 2019).

It is however worth noting that in conditions inducing N_2_ fixation, filamentous diazotrophic cyanobacteria undergo a major metabolic readjustment that involves also the photosynthetic apparatus. Photosynthetic components are segregated between VCs and HCs indeed (Cardona *et al*., 2009). In HCs, PSII is dismantled to avoid water splitting and preserve a microoxic environment, whereas PSI is the only photosystem left active (Magnuson & Cardona, 2016). PSII-independent electron pathways are therefore expected also to contribute to the generation of the proton gradient across the thylakoid membrane in HCs (Zheng *et al*., 2019; Bandyopadhyay *et al*., 2021) and fulfil the generation of ATP via photophosphorylation (Kumar *et al*., 2010; Magnuson, 2019; Bandyopadhyay *et al*., 2021) to fuel the nitrogenase activity (Cardona & Magnuson, 2010).

In this work we interestingly observed that upon high light and excess CO_2_ the efficiency of CEF exceeded LEF, and the former represented the major contribution to the total photosynthetic activity indeed (Fig. 5b and 5C). In this condition, growth also reached the maximal value observed (Fig. 1b) and the correlation analysis showed that it did correlate positively with the cyclic electron flow around PSI (Fig. 6a).

The cyclic electron fluxes around PSI are known to contribute to the generation of ATP indeed. Two complementary pathways have been identified in cyanobacteria, involving the molecular complexes NDH and PGR5/PGRL1 (Mullineaux, 2014). Both contribute to the generation of the proton motive force that sustains the synthesis of ATP (Allahverdiyeva *et al*., 2013) but the pathway mediated by the NDH complex is suggested to contribute the most (Miller *et al*., 2021) through the translocation of up to 4 protons per electron across the thylakoid membrane (Battchikova *et al*., 2011; Peltier *et al*., 2016). The cyclic electron transport is suggested to have a basal energetic contribution, to cover for the extra chemical energy needed for C fixation via the CBB cycle in all photosynthetic organisms, that the linear electron flux from PSII cannot sustain alone (Theune *et al*., 2021). The ATP/NADPH ratio to fix one molecule of substrate is greater for the nitrogenase (16 ATP and 8 NADPH) than the CBB cycle (3 ATP and 2 NADPH), i.e. 2 vs 1.5, respectively, reasonably indicating that the contribution of the PSII-independent photosynthetic activity towards the energy balance of the host is more relevant in diazotrophic cyanobacteria upon N_2_ fixing conditions, likely to fulfil the large energetic demand of the N_2_ fixation reaction.

In this work, the cyclic electron flow around PSI significantly increased upon diazotrophic with respect to NO_3_^-^ replete conditions indeed (Fig. 5a), supporting this hypothesis.

Overall, these observations suggest that, upon diazotrophic conditions, the PSII-independent photosynthetic activity has a predominant role towards the energy balance of the host rather than photoprotection. The establishment of the N_2_ fixation capacity likely compensates for the reduced contribution of CEF towards photoprotection, whilst preserving the ability to withstand high light.

### Photosynthesis regulation likely contributes to the functional coordination between cell types upon diazotrophic conditions

In diazotrophic conditions and atmospheric CO_2_, photosynthesis was fully active (Fig. 4b) and the PSII-dependent activity of VCs represented the major contribution (Fig. 5b). Because the CBB cycle was here limited by the CO_2_ availability, most of the photosynthetic activity is likely used to feeding the N_2_ fixation reaction (Fig. 7a and 7b), overall suggesting that in this condition the functional coordination between the two cell types is unbalanced towards HCs (Fig. 7a).

**Fig. 7:**
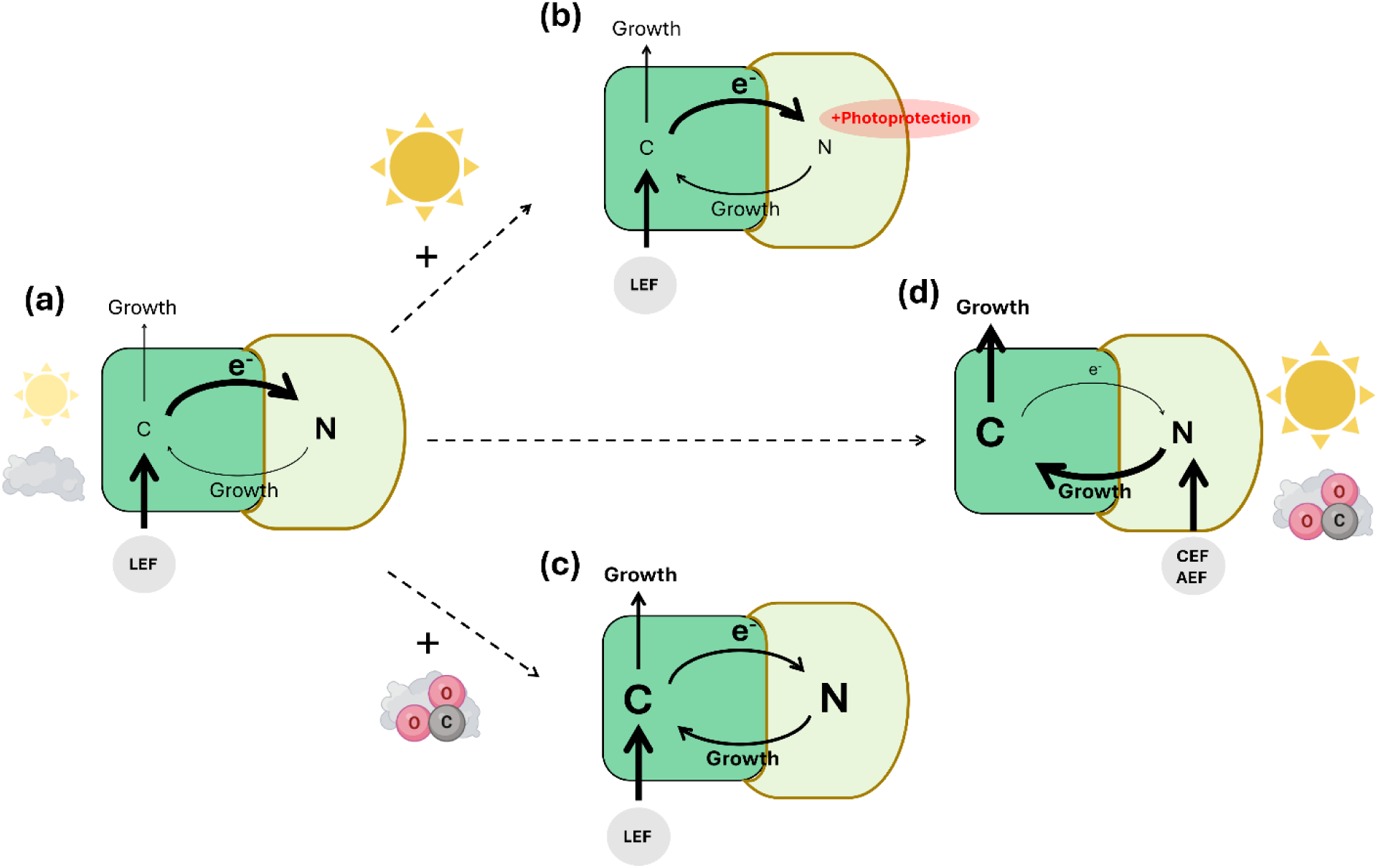
Model of intercellular functional coordination in *Anabaena*. Schematic representation showing the contribution of different electron fluxes of photosynthesis on the regulation of the intercellular functional coordination between C and N assimilation pathways, according to changing environmental conditions. The suggested preferential coordination routes upon atmospheric (a and b) or excess CO_2_ (c and d) and low (a and c) or high light (b and d) are highlighted by the thickness of the arrows. Green, vegetative cells; pale green, heterocysts. LEF, PSII-dependent photosynthetic activity specific of VCs; CEF and AEF, PSII-independent photosynthetic activity shared between VCs and HCs.

Upon light excess (Table 1), the PSII-dependent photosynthetic activity still represented the major contribution to TEF (Fig. 5b). In this condition the functional coordination between cell types is therefore still likely unbalanced towards HCs but to exploit the large electron sink capacity of nitrogenase as relief valve to prevent the overreduction of the photosynthetic transport chain and withstand light saturation (Fig. 7b).

On the other hand, upon higher CO_2_ supply the CBB cycle increased its electron sink capacity. In low light, the PSII-dependent photosynthesis still represented the major contribution to TEF (Fig. 5b), suggesting that the photosynthetic activity of VCs is here contributing to sustain both the CBB cycle and the nitrogenase complex in HCs (Fig. 7c). Therefore, the functional coordination between the two cell types is here likely more balanced (Fig. 7c).

Upon higher light, the mutant strain showed the greatest reduction of growth with respect to the WT (Fig. 1e), suggesting that the flux of fixed N towards VCs *via* cyanophycin provides the largest contribution on growth among the conditions tested. Therefore, we suggest that the functional coordination between the two cell types is here unbalanced towards VCs (Fig. 7d). Interestingly, the major contribution to TEF was provided by the cyclic electron flow around PSI, that indeed exceeded the efficiency of LEF (Fig. 5c). Moreover, in this condition photosynthesis was likely not light saturated as the CBB cycle operated upon excess CO_2_ and the nitrogenase further expanded the total electron sink capacity of the host, suggesting a major contribution of CEF towards the energy balance of the host.

Overall, this work suggests that in diazotrophic cyanobacteria, the PSII-independent photosynthetic activity is interestingly placed at the intersection between the regulation of photosynthesis to withstand excess irradiation and the feeding of the N_2_ fixation reaction, highlighting a likely active role also on the regulation of the functional coordination between the metabolism of C and N upon diazotrophic conditions.

## Acknowledgements

F.A. gratefully acknowledges support from the Italian Ministry of University and Research (Rita Levi Montalcini Program) and the Fondazione Cariparo (Starting Package Program). G.P. acknowledges the support from the University of Padova STARS-Stg Project “WWBIOMASS”, Center “Giorgio Levi Cases” Project “REENACT” and the Department of Biology BIRD2023 “ORGANISM”.

## Author contributions

AS, experimental set-up, data collection and visualization, drafting original manuscript; MS, experimental set-up and data collection; MM, statistical analysis; AF, data collection for the nitrogen fixation assay; FA, protocol set-up for the nitrogen fixation assay; EF, contributed strain CSMI6 and provided critical revision and editing of the manuscript; TM, critical revision and editing of the manuscript; GP, conception and design, writing, revision and editing of the manuscript.

## Data availability

The authors confirm that the data supporting the findings of this study are available within the article or its supplementary materials.

