## Supplementary Information for "Photosynthesis regulation impacts carbon and nitrogen assimilation in the diazotrophic cyanobacterium *Anabaena* sp. PCC 7120"

### Supplementary Figures

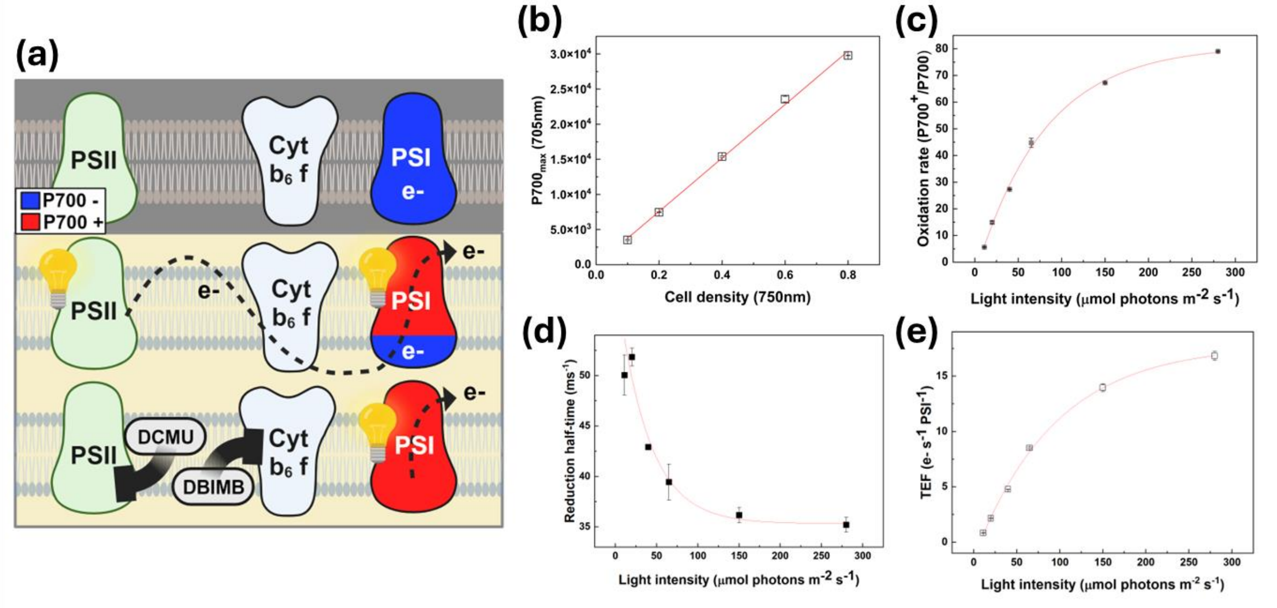


**Supplementary Fig. S1:** Estimation of PSI content and photosynthetic activity in *Anabaena* using JTS-10 absorption spectroscopy. (a) Schematic representation of the technique. In the dark (dark background), PSI is completely reduced (blue). When exposed to light (light background) PSI is oxidized by light and re-reduced by the electron flux coming from PSII. Inhibitors (i.e. DCMU or DBIMB) selectively block the electron fluxes toward PSI, enabling its complete oxidation upon light exposure. (b) Maximum 705 nm absorption ($P700_{max}$) of *Anabaena* cultures at different cell concentrations ($OD_{750nm}$), upon exposure to saturating light in presence of PSII and Cyt $b_6f$ inhibitors. (c-e) Parameters calculated from P700 traces of *Anabaena* cultures upon exposure to increasing light intensities (11, 20, 40, 65,150, 280 µmol photons $m^{-2}$ $s^{-1}$): (c) oxidation rate ($P700^+$/P700) upon illumination; (d) reduction half-time (Reduction rate) in light-dark transition; (e) calculated TEF for each given light intensity. In C, D and E data represents averages of at least three biological replicates (± SD).

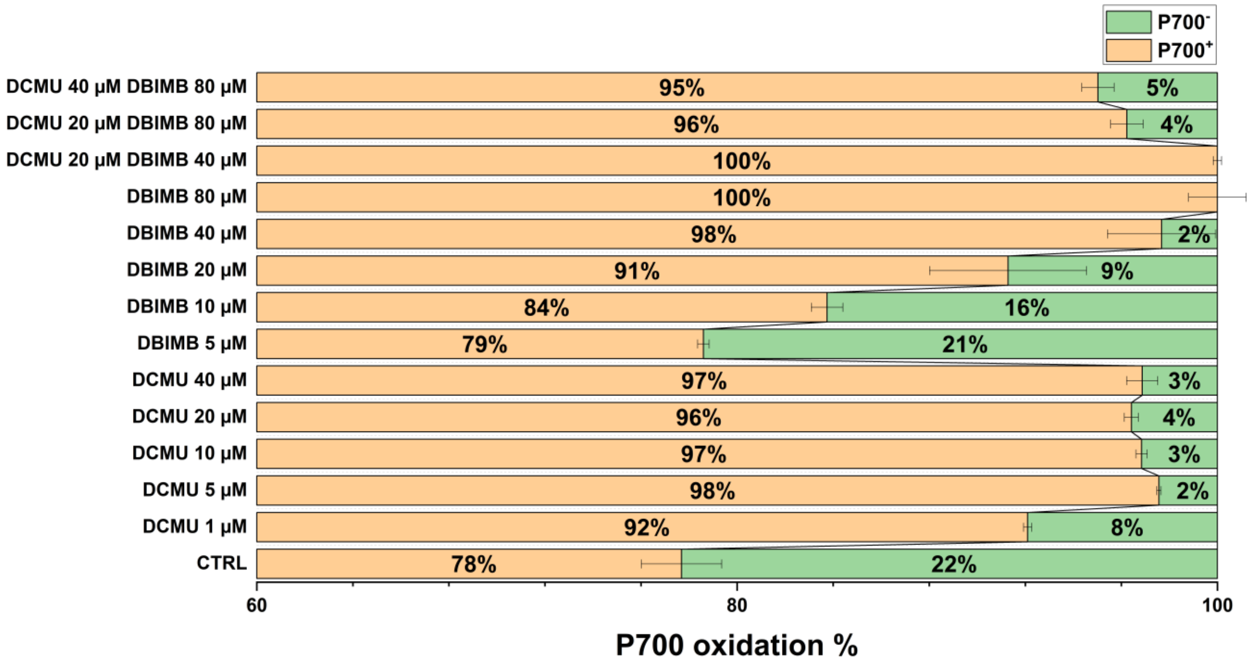


**Supplementary Fig. S2:** Dose evaluation of PSII and Cyt $b_6f$ inhibitors. The effectiveness of inhibitors was evaluated by 705 nm absorption during saturating light exposure (2080 µmol photons $m^{-2}$ $s^{-1}$). DCMU blocks the PSII-dependent electron flow, resulting in an almost complete oxidation of PSI upon illumination. Maximum absorption was observed above a 5 µM dose of the inhibitor. In combination with DCMU, DBIMB blocks the alternative electron flux to PSI, so to enable its complete oxidation. Maximum 705 nm absorption was observed in co-presence of 20 µM DCMU and 40 µM DBIMB. Higer doses caused a reduction of the signal likely due to side effects of the inhibitors (i.e. photoinhibition).

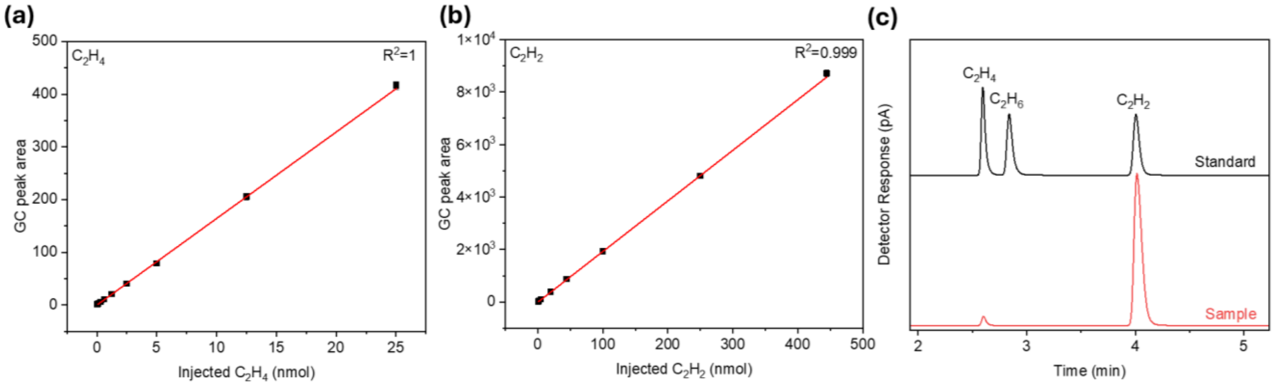


**Supplementary Fig. S3:** Chromatography quantification of ethylene. Calibration curves for gas chromatography quantification of ethylene (a) and acetylene (b) using GC-TCD-FID with the corresponding coefficients of linear correlation ($R^2$). (c) Typical gas chromatograms for the reduction reaction of acetylene to ethylene. Gas chromatograms (retention time of $C_2H_4$, $C_2H_6$, $C_2H_2$) of the $C_2H_4$ standard (2.5 vol.%), $C_2H_6$ standard (2.5 vol.%), $C_2H_2$ standard (2.5 vol.%) (black), and the headspace composition of our system under $C_2H_2$ ($\geq$ 99.5 vol.%) (red).

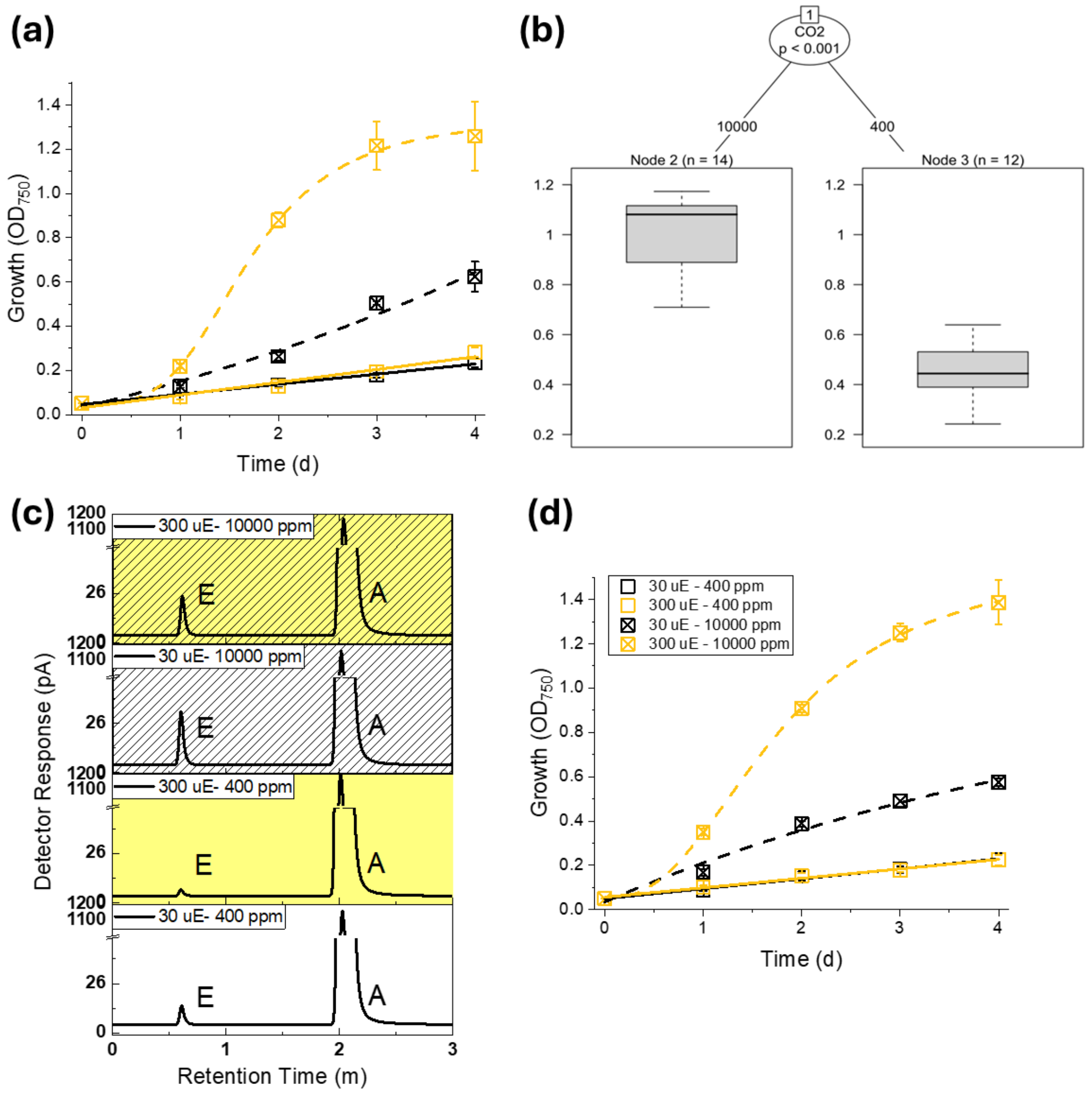


**Supplementary Fig. S4:** Growth and $N_2$ fixation of *Anabaena* upon different light and $CO_2$ conditions. (a) Growth, expressed as $OD_{750}$, upon $NO_3^-$ replete conditions for *Anabaena* WT, upon different $CO_2$ and light availability. (b) Partitioning tree of the growth rate data for *Anabaena* WT. The impact on the growth rate of *Anabaena* of the different experimental variables was investigated altogether, using the formula Growth rate ~ $CO_2$ availability + Number of photons + N availability. After Bonferroni correction, only $CO_2$ availability was observed to significantly impact the growth rate, dominating on the marginal effects of light and $NO_3^-$ availability. (c) Gas chromatograms for the reduction reaction of acetylene (A) to ethylene (E) upon different $CO_2$ and light availability in diazotrophic conditions for *Anabaena* WT. (d) Growth, expressed as $OD_{750}$, upon $NO_3^-$ replete conditions for *Anabaena* mutant CSMI6, upon different $CO_2$ and light availability. µE = µmol photons $m^{-2}$ $s^{-1}$, ppm = ppm of $CO_2$. Data represents averages of > three biological replicates (± SD).

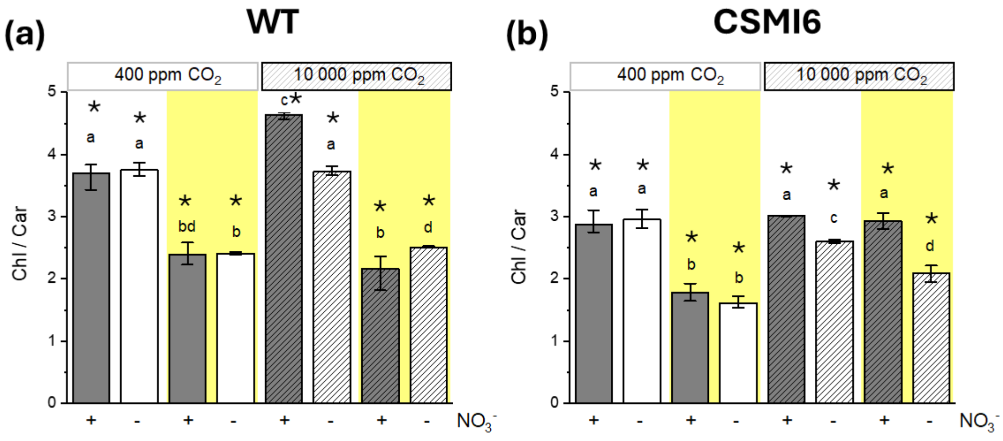


**Supplementary Fig. S5:** Chlorophyll to Carotenoids ratio upon different light and $CO_2$ conditions. (a-b) Chl/Car ratio upon different irradiances and $CO_2$ availability for *Anabaena* WT (a) and for *Anabaena* mutant CSMI6 (b), comparing $NO_3^-$ replete with diazotrophic conditions. Grey and empty bars indicate $NO_3^-$ replete and diazotrophic conditions, respectively; white and yellow backgrounds indicate limiting (30 µmol photons $m^{-2}$ $s^{-1}$) or excess (300 µmol photons $m^{-2}$ $s^{-1}$) irradiance, respectively; full colours and fill pattern indicate limiting (400 ppm) or excess (10000 ppm) $CO_2$, respectively. Data represents averages of at least three biological replicates (± SD). Different lowercase letters and asterisks indicate statistically significant differences between samples upon different environmental conditions within the same strain and between strains upon the same environmental condition, respectively (one-way ANOVA, $p < 0.05$).

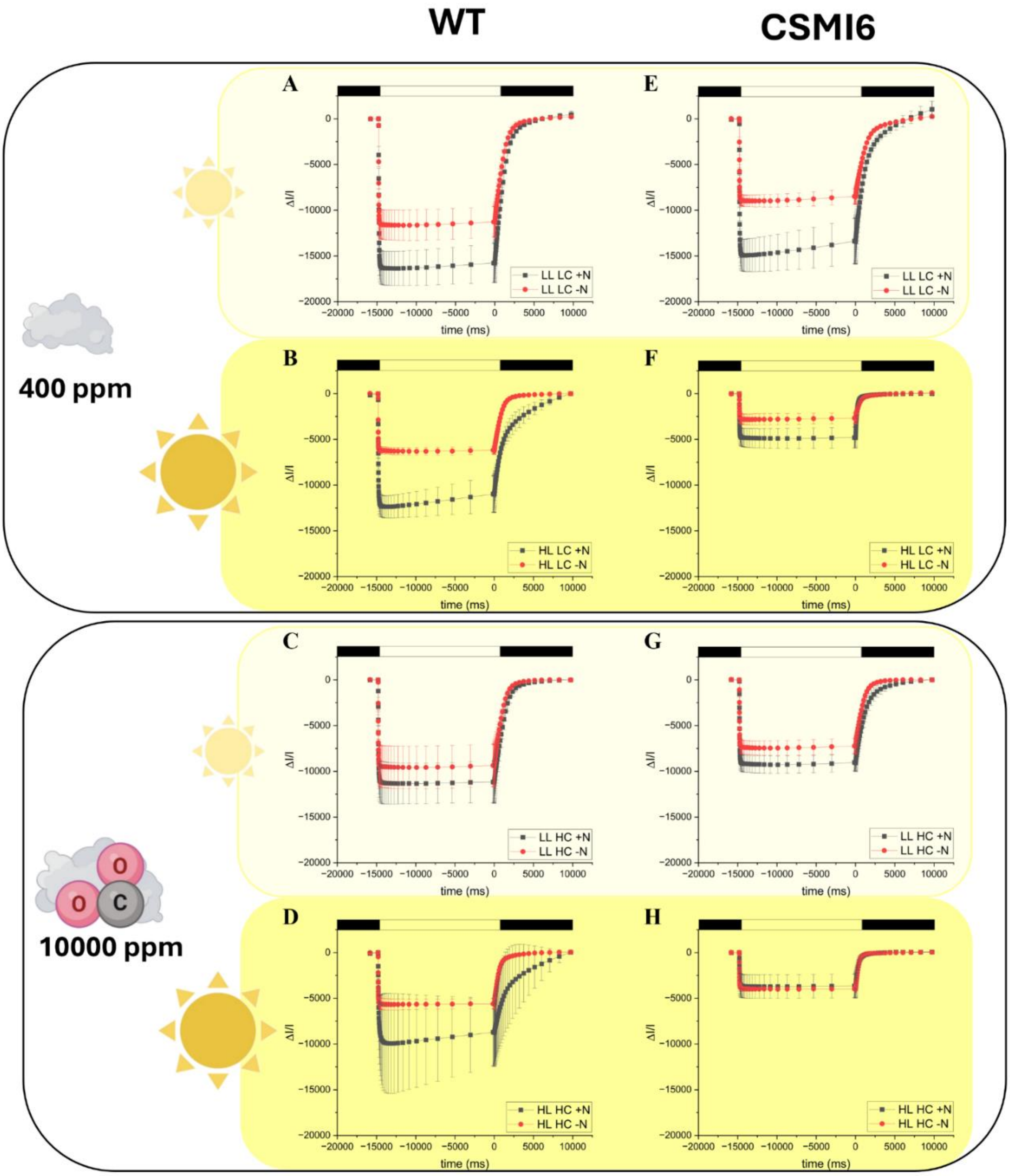


**Supplementary Fig. S6:** Representative traces for the PSI quantification in *Anabaena*. The PSI content upon the different irradiances and $CO_2$ availability tested in this work was evaluated from the maximum absorption of $P700^+$ at 705 nm, in the presence of DCMU and DBMIB and a saturating light of 2080 µmol photons $m^{-2}$ $s^{-1}$ (see methods for details), comparing $NO_3^-$ replete (black) with diazotrophic (red) conditions. (a, b, c, d) Data for *Anabaena* WT, whilst (e, f, g, h) data for *Anabaena* CSMI6. (a, e, b, f) Data upon exposure to atmospheric $CO_2$, whilst (c, d, g, h) data upon exposure to excess $CO_2$. Light and dark yellow backgrounds indicate exposure to low and excess light, respectively. White box, actinic light on; black box, actinic light off. Data represents averages of at least three biological replicates (± SD).

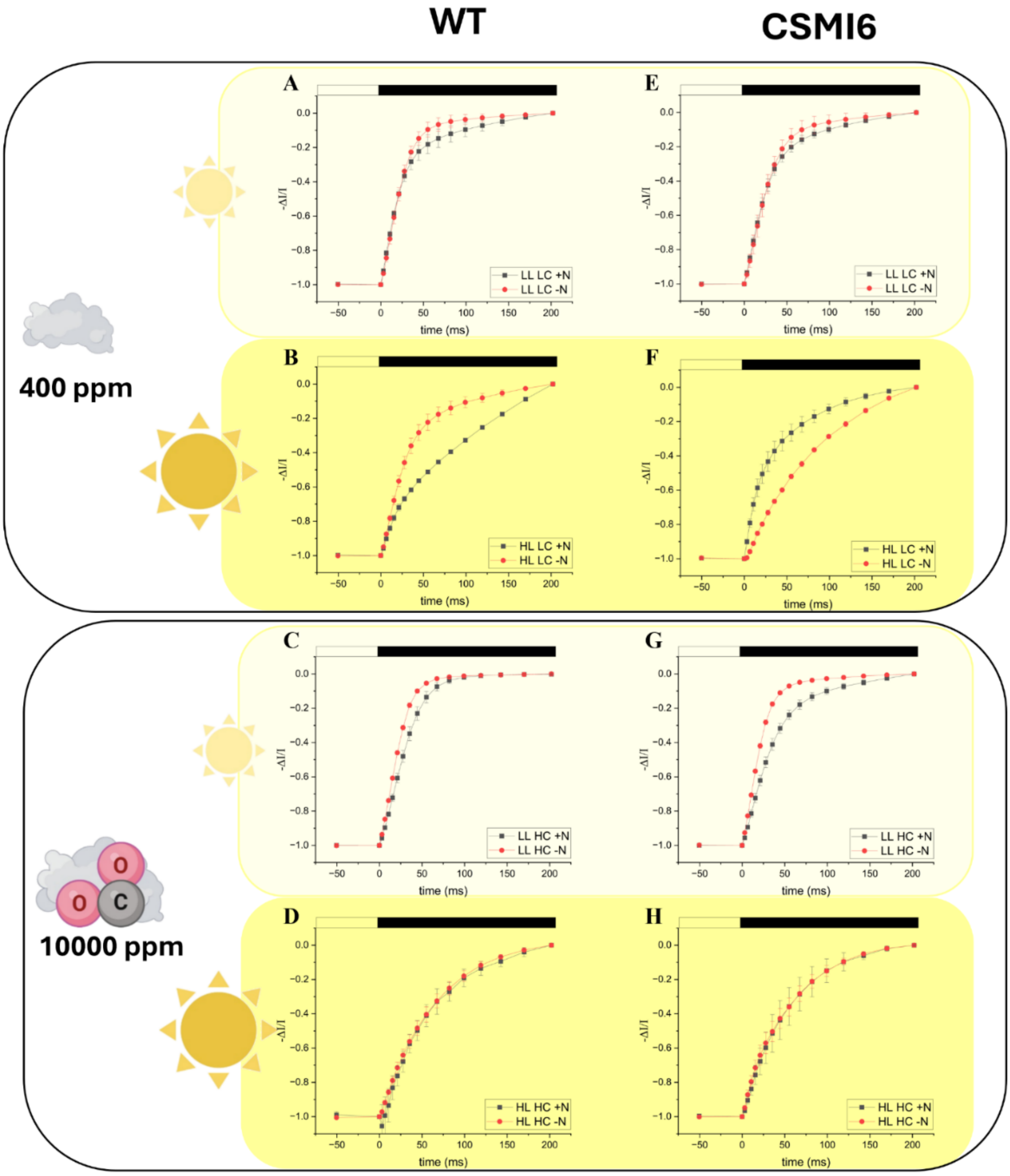


**Supplementary Fig. S7:** Representative traces for the quantification of the Total Electron Flow in *Anabaena*. The Total Electron Flow (TEF) per unit of PSI and time ($s^{-1}$) upon the different irradiances and $CO_2$ availability tested in this work was evaluated from the re-reduction kinetics of oxidized $P700^+$. Quantification was performed upon exposure to a saturating light intensity (2080 μmol photons $m^{-2}$ $s^{-1}$) before evaluating the recovery kinetics in the dark, here reported, comparing $NO_3^-$ replete (black) with diazotrophic (red) conditions. (a, b, c, d) Data for *Anabaena* WT, whilst (e, f, g, h) data for *Anabaena* CSMI6. (a, e, b, f) Data upon exposure to atmospheric $CO_2$, whilst (c, d, g, h) data upon exposure to excess $CO_2$. Light and dark yellow backgrounds indicate exposure to low and excess light, respectively. White box, actinic light on; black box, actinic light off. Data represents averages of at least three biological replicates (± SD).

**Supplementary Tables**

**Supplementary Table S1:** Analysis of variance. Physiological parameters measured in both the parental (wild-type) and mutant strain (CSMI6) were correlated. In bold the parameters changing the most between the two strains, regardless of any environmental condition tested in this work. TEF, Total Electron Flow; LEF, Linear Electron Flow; CEF, Cyclic Electron Flow; AEF, Alternative Electron Flow; P700, PSI content; Chl, Chlorophyll content; Car, Carotenoid content.

| Parameters | Ratio Variance of CSMI6/wild-type | *p* value |
|---|---|---|
| Growth rate ($d^{-1}$) | 1.061813 | 0.4419 |
| TEF ($e^{-}$ $s^{-1}$ $PSI^{-1}$) | 0.8 | 0.7 |
| **AEF ($e^{-}$ $s^{-1}$ $PSI^{-1}$)** | **4.03** | **0.0003904** |
| LEF ($e^{-}$ $s^{-1}$ $PSI^{-1}$) | 1.21 | 0.3158 |
| CEF ($e^{-}$ $s^{-1}$ $PSI^{-1}$) | 0.95 | 0.54 |
| P700 | 1.15 | 0.359 |
| **Chl (µg $mg^{-1}$)** | **2.649715** | **0.008336** |
| **Car (µg $mg^{-1}$)** | **3.627026** | **0.0009027** |
